# Stabilising effects of competition and diversity determine vaccine impact on antibiotic resistance evolution

**DOI:** 10.1101/610188

**Authors:** Nicholas G. Davies, Stefan Flasche, Mark Jit, Katherine E. Atkins

## Abstract

Bacterial vaccines can protect recipients from contracting potentially antibiotic-resistant infections. But by altering the selective balance between sensitive and resistant strains, vaccines may also suppress—or spread—antibiotic resistance among unvaccinated individuals. Predicting the outcome requires knowing what drives selection for resistance in bacterial pathogens, and in particular, what maintains the circulation of both antibiotic-sensitive and resistant strains of bacteria. Using mathematical modelling, we show that the frequency of penicillin resistance in *Streptococcus pneumoniae* (pneumococcus) across 27 European countries can be explained by between-host diversity in antibiotic use, heritable diversity in pneumococcal carriage duration, or frequency-dependent selection brought about by within-host competition between resistant and sensitive strains. We use our calibrated models to predict the impact of non-serotype-specific pneumococcal vaccination upon the prevalence of carriage, incidence of disease, and frequency of resistance for *S. pneumoniae*. We find that the relative strength and directionality of competition between resistant and sensitive pneumococcal strains is the most important determinant of whether vaccination promotes, inhibits, or has little effect upon the evolution of antibiotic resistance. Finally, we show that country-specific differences in pathogen transmission substantially alter the predicted impact of vaccination, highlighting that policies for managing resistance with vaccines must be tailored to a specific pathogen and setting.

**One sentence summary:** Frequency-dependent competition and extrinsically-maintained diversity shape selection for antibiotic resistance following vaccination.

---

In an age of widespread antibiotic resistance, there is growing interest in using vaccines to prevent bacterial infections that would otherwise call for treatment with antibiotics (*1-4*). This interest arises for two main reasons: first, vaccines are effective against both antibiotic-resistant and antibiotic-sensitive bacteria; and second, successful prophylaxis removes the need for a course of antibiotic therapy that might promote more resistance (*2–5*). Over the past two decades, the use of pneumococcal conjugate vaccines (PCVs) has seemingly borne out these advantages. Administering PCVs to young children has substantially reduced disease caused by *S. pneumoniae (5–8*)—a common asymptomatic coloniser of the nasopharynx which can cause pneumonia, meningitis and other infections when invasive—and has decreased demand for antibiotic therapy, largely by reducing cases of otitis media (*5, 9*). But because PCV formulations target only a fraction of the ~100 known pneumococcal serotypes, the niche vacated by PCV-targeted serotypes has been filled by non-vaccine serotypes, and overall pneumococcal carriage has rebounded to pre-vaccine levels (*10, 11*). Concomitantly, the incidence of infections attributed to non-vaccine serotypes (*12*) and the proportion of non-vaccine-type infections exhibiting antibiotic resistance (*5, 13*) have risen in many settings. Concern over serotype replacement—along with the high cost of PCV manufacturing—has spurred the development of “universal” (non-serotype-specific) whole-cell or protein-based pneumococcal vaccines protecting against all serotypes, some of which are now in early-stage clinical trials (*14*). If successful, universal pneumococcal vaccines could reduce the burden of pneumococcal disease without selecting for serotype replacement.

However, it is unclear how universal vaccination itself may impact upon the evolution of antibiotic resistance in *S. pneumoniae*, which is a concern given that vaccination is unlikely to eliminate pneumococcal carriage entirely (*15*). Mathematical models of bacterial transmission can be used to predict the impact of vaccination on antibiotic resistance (*16, 17)*, but existing models for *S. pneumoniae* focus on serotype-specific vaccines and, even then, disagree over the expected impact of vaccination on resistance evolution (*18–24*). Comparing and interpreting the results of these models is hampered by the fact that none starts from a position of recapitulating large-scale empirical patterns of antibiotic resistance. The main challenge in replicating these patterns lies in identifying the mechanisms that maintain long-term coexistence between sensitive and resistant pneumococcal strains across a wide range of antibiotic treatment rates, like those seen across Europe and the United States (*25, 26*). Robust predictions of the long-term impact of non-serotype-specific vaccination on resistant pneumococcal disease require a mechanistic understanding of these patterns.

## Results

### Stability in resistance evolution can be maintained by frequency-dependent competition or extrinsically-imposed diversity

A model must be able to explain the current burden of an infectious disease before it can be used to robustly predict the impact of interventions for managing that disease. Across Europe, the frequency of antibiotic resistance among isolates from pneumococcal infections shows two salient features for models to recapitulate (Fig. S1). One feature is spatial: the frequency of penicillin non-susceptibility varies between countries, and is higher in countries where more penicillin is consumed (*27*). The other is temporal: although in individual countries, resistance fluctuates from year to year, the overall frequency across Europe of penicillin non-susceptibility in pneumococcal isolates has remained steady at roughly 12% since consolidated records began in 2005 (*28*). These observations contradict simple models of resistance evolution, which predict that intermediate frequencies of resistance cannot be stably maintained in the long term: that is, either sensitive strains will competitively exclude resistant strains, or resistant strains will competitively exclude sensitive strains, unless there is some mechanism that maintains coexistence between them (*25, 29*).

By conducting a literature search, we identified nine such mechanisms (*25, 26, 30–41*) that fall into two broad classes. In one class, coexistence is maintained by environmental or genetic diversity that effectively creates separate niches for resistant and sensitive strains, preventing them from completely overlapping in competition. In the other class, competition between resistant and sensitive strains is itself the stabilising factor that maintains coexistence, because resistant and sensitive strains exhibit alternative competitive phenotypes that afford strains a competitive advantage when rare, thus promoting negative frequency-dependent selection for resistance. Thus, extrinsically-imposed diversity and frequency-dependent competition are two key forces maintaining stability in resistance evolution. We find that four of the nine identified mechanisms for maintaining coexistence are biologically plausible for *S. pneumoniae* (Table 1).

**Table 1.** Mechanisms for maintaining coexistence.

|  | Mechanism | Mode of action | Plausible mechanism for coexistence in *S. pneumoniae*? |  | Consistent with empirical patterns? (Fig. 2) |  |
| --- | --- | --- | --- | --- | --- | --- |
| DIVERSITY | Treatment diversity | Assortatively-mixing subpopulations differ in treatment rates (25, 34–37) | ✓ | Yes | ✓ | Yes |
|  | Pathogen diversity | Subtypes maintained by diversifying selection differ in propensity for resistance (38) | ✓ | Yes | ✓ | Yes |
|  | Treated class | Individuals currently in treatment maintain resistant strains (25, 34, 39, 40) | ✗ | No: Only supports a small amount of coexistence (25) | ■ | N/A |
|  | Within-host niches | Sensitive and resistant strains exploit separate niches within the host (30, 41) | ✗ | No: Resistant and sensitive strains are known to occupy the same niches (29) | ■ | N/A |
|  | Mutation pressure | Mutation-selection balance maintains intermediate resistance frequency (30, 31, 37) | ✗ | No: De novo acquisition of resistance in *S. pneumoniae* is not frequent enough (25) | ■ | N/A |
|  | Prescription feedback | Doctors reduce prescribing of a drug as resistance to it increases (37, 39) | ✗ | No: Does not explain how coexistence is maintained over a range of treatment rates | ■ | N/A |
| COMPETITION | Within-host competition: Treatment competition | Within-host competition creates frequency-dependent selection for resistance (25, 26, 32, 33, 40) | ✓ | Yes | ✓ | Yes |
|  | Within-host competition: Growth competition | “ | ✓ | Yes | ✓ | Yes |
|  | Superinfection | Superinfection creates frequency-dependent selection for resistance (30) | ✗ | No: Requires resistant strain to transmit better than sensitive strain in absence of antibiotics | ■ | N/A |

### Four models of resistance evolution

To compare these four mechanisms, we embed each in a shared model framework of person-to-person transmission of nasopharyngeal pneumococcal carriage. This framework tracks the country-specific frequency of resistance in pneumococci circulating among children under five years old, the age group that drives the majority of pneumococcal transmission and disease (*42, 43*). We assume that each individual makes effective contact with another random individual at rate β, thereby potentially acquiring a strain (either sensitive or resistant) carried by the contacted person. With probability *c*, resistant strains fail to transmit, where *c* represents the transmission cost of resistance (*44, 45*). A carrier naturally clears all strains at rate *u*, and is exposed to antibiotic therapy at a country-specific rate *τ*, which clears the host of sensitive strains only. We assume this treatment rate is independent of carriage status (*46*) and we do not explicitly track disease progression in hosts.

Under the “Treatment diversity” and “Pathogen diversity” models, extrinsically-maintained diversity among hosts or among pathogens prevents competitive exclusion by keeping resistant and sensitive strains from fully competing with each other. In the “Treatment diversity” model (Fig. 1a), heterogeneity in the consumption of antibiotics between host subpopulations within a country maintains coexistence (*25, 34, 35*). These subpopulations could correspond to geographical regions, socioeconomic strata, host age and risk classes, or a combination of these. Provided that transmission between high-consumption (resistance-promoting) and low-consumption (resistance-inhibiting) subpopulations is not too frequent, an intermediate frequency of resistance can be maintained across the whole population. Because coexistence is maintained by assortative mixing between subpopulations differing in antibiotic use, the key parameters governing coexistence in this model are κ, the variability in antibiotic consumption between subpopulations, and *g*, the relative rate at which within-country contact is made within subpopulations rather than between them (Fig. S2).

**Fig. 1.**
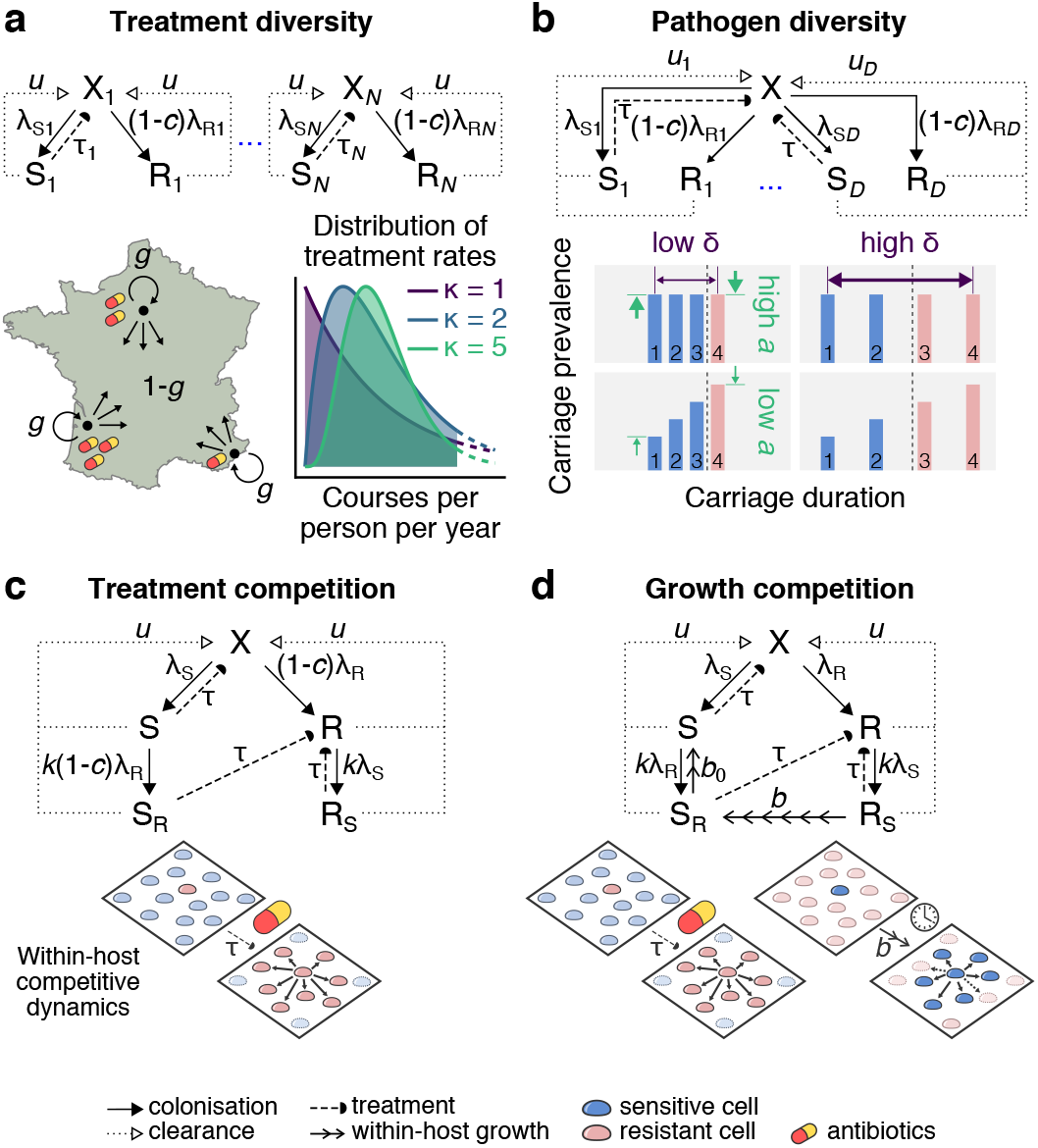
Four models of resistance evolution. X hosts are uncolonised, S hosts are colonised with the sensitive strain and R hosts are colonised with the resistant strain. Force of infection terms *λ_A_* are equal to the person-to-person contact rate β times the probability that a contacted individual carries strain *A; c* is the transmission cost of resistance; *u* is the natural clearance rate; and *τ* is the rate of antibiotic treatment. **(a)** “Treatment diversity”: each country is split into subpopulations varying in treatment rate *τ_i_*, with treatment rates drawn from a gamma distribution with shape *κ*. Within a country, individuals assort with their own subpopulation with probability *g;* this assortative mixing among treatment-varying subpopulations allows coexistence between sensitive and resistant strains. **(b)** “Pathogen diversity”: the pathogen comes in multiple subtypes maintained by diversifying selection, each with its own natural carriage duration *u*_*d*^−1^_. Diversifying selection is stronger *(i.e.*, more equalizing) as *a* increases, while carriage durations span a greater range as δ increases. Only those subtypes whose carriage duration exceeds a critical threshold (dashed line) are selected for resistance, so that overall, both sensitive and resistant strains can circulate. **(c)** “Treatment competition”: singly-colonised hosts can acquire a small amount of another strain at relative rate *k* (host states S_R_ and R_S_). Dually-colonized hosts only transmit the dominant strain, so there is within-host competition between co-colonising strains. Population-level coexistence is maintained by treatment-mediated within-host competition. **(d)** “Growth competition”: as in panel c, but the transmission cost of resistance is removed and sensitive strains now outgrow resistant strains within co-colonised hosts at rate *b.* Coexistence is maintained by both treatment-mediated and growth-mediated within-host competition. Panels a-d illustrate alternative model dynamics for a single country; our full model framework tracks dynamics for 27 European countries simultaneously, which themselves mix with assortativity *f.*

In the “Pathogen diversity” model (Fig. 1b), pneumococci are divided into subtypes (“D-types”(*38*)) that vary in their mean duration of natural carriage. All else equal, the D-type with the longest carriage duration would be expected to competitively exclude all other strains; the model assumes that diversifying selection acting on the D-type locus keeps all subtypes in circulation. What D-types correspond to is not explicitly specified by this model, but one candidate is serotype variation. For example, if antigenic diversity is promoted by host acquired immunity to capsular serotypes, and serotypes tend to differ in their intrinsic ability to evade clearance by the immune system, then intermediate resistance can be maintained because selection for resistance tends to be greater in serotypes that have a longer duration of carriage (*38*). Long-lasting serotypes will tend to evolve resistance, while shorter-lived serotypes will tend not to—a pattern observed in *S. pneumoniae (38*) and reproduced by this model (Fig. S3). The parameters governing coexistence in this model are *a*, the strength of diversifying selection on the D-type locus, and δ, the variability between subtypes in clearance rate.

Under the “Treatment competition” and “Growth competition” models, coexistence is maintained by within-host competition between sensitive and resistant strains. In these models, hosts can be co-colonised by multiple strains. Then, competition between strains within the host niche determines which strain is transmitted to other potential hosts (*26*). The “Treatment competition” model (Fig. 1c) assumes that antibiotic therapy mediates within-host competition, such that when a co-colonised host takes antibiotics *(i.e.*, at rate τ), the sensitive strains are cleared and only the resistant strains are transmitted to other hosts. The “Growth competition” model (Fig. 1d) has both treatment-mediated and growth-mediated competition: while in the presence of antibiotics, resistant strains still outcompete co-colonising sensitive strains, in the absence of antibiotics, sensitive strains gradually outcompete co-colonising resistant strains at rate *b.* We assume that there is no transmission cost of resistance in this latter model *(i.e., c* = 0); instead, the within-host growth advantage *b* of sensitive strains accounts for the cost of resistance. In these competition models, resistant strains have an advantage in antibiotic-mediated competition, while sensitive strains have an advantage in growth-mediated competition. These alternative forms of within-host competition can both promote coexistence because rare strains can more consistently exploit a competitive advantage over common strains, thus creating negative frequency-dependent selection for resistance (*26*). The key parameter governing coexistence in these two models is *k*, the relative rate of co-colonisation compared to primary colonisation.

In all four models, we assume that contact between individuals is assortative by country, such that with probability *f*, contact is with a random person from the same country, and with probability 1 - *f*, contact is with a random person from any country. We implement these models using systems of ordinary differential equations. All four models (*25, 26, 38*) are structurally neutral (*25, 29*), meaning that any coexistence exhibited by the models is accounted for by the specified biological mechanism rather than by any bias in the logical structure of the model that generates coexistence “for free” (*29*). Additionally, while the within-host competition models capture co-colonisation using a simplified subset of only 2 “mixed-carriage” states (S_R_ and R_S_, Fig. 1a&b), we have previously shown (*26*) that this is equivalent to a more complex individual-based model with an arbitrary number of mixed-carriage states.

### All four models reproduce observed patterns of resistance

The European Centre for Disease Prevention and Control (ECDC) monitors antibiotic consumption and resistance evolution across European countries (*13, 28*). These data capture a natural experiment in resistance evolution: for each monitored drug and pathogen, each country reports a different rate of antibiotic consumption in the community and exhibits a different frequency of resistance among invasive bacterial isolates. By fitting models to this multi-country data set, we can potentially rule out models that cannot reproduce the large-scale patterns that are observed. We use Bayesian inference to fit the model-predicted equilibrium frequency of resistance to the reported frequency of penicillin non-susceptibility in *S. pneumoniae* across 27 European countries, assuming a 50% carriage prevalence (*11, 42*) and a carriage duration of 47 days (*47, 48*) in children under five years old. We begin by assuming that countries only differ by their reported treatment rate—where we define a treatment course as equivalent to *z* = 5 defined daily doses of penicillin—with other model parameters shared across countries.

Strikingly, each model fits equally well to the empirical relationship between resistance and antibiotic use (all model WAICs are similar; Fig. 2a) and recovers plausible posterior parameter distributions (Fig. 2b; Fig. S4). That is, the empirical data do not distinguish between the four alternative mechanisms of resistance evolution we have identified. Later, we relax the assumption that only the treatment rate varies between countries, allowing us to capture additional between-country variation in resistance not explained by population-wide penicillin consumption.

**Fig 2.**
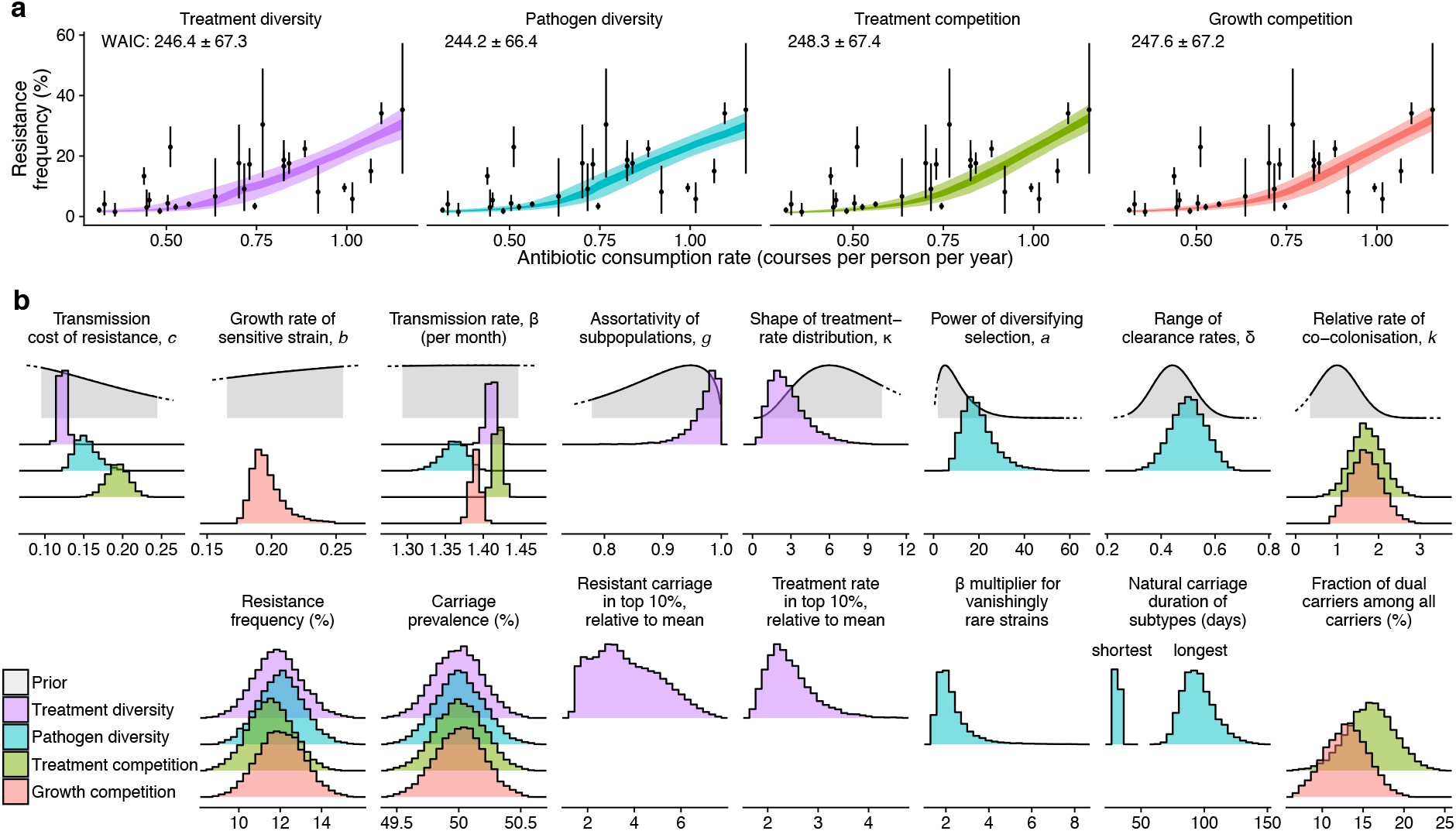
Four models reproduce patterns of resistance in *S. pneumoniae* in Europe. **(a)** Model fits with associated WAIC (± standard error). Points and vertical lines show the mean and 95% highest density intervals (HDIs) for the reported proportion of invasive *S. pneumoniae* isolates that are resistant to penicillin plotted against the penicillin consumption rate in under-5s. Ribbons show the 50% and 95% HDIs for resistance prevalence from each fitted model. **(b)** The top row shows estimated posterior distributions for the free parameters in each model; the bottom row shows model outputs associated with these parameters to aid interpretation.

### Mechanisms of resistance evolution determine the impact of vaccination on resistant disease

To determine the impact of universal vaccination on pneumococcal disease, we consider three outcomes. The first is the impact of the vaccine upon the prevalence of pneumococcal carriage. The second is the vaccine impact upon the frequency of penicillin resistance among circulating pneumococcal strains remaining after vaccination. The third is the impact of the vaccine upon the prevalence of resistant pneumococcal carriage—*i.e.*, the prevalence of carriage multiplied by the frequency of penicillin resistance. Since all four models are equally capable of recapitulating observed patterns of penicillin resistance in *S. pneumoniae*, our aim is to determine whether the mechanism maintaining stability in resistance evolution—frequency-dependent competition or extrinsically-imposed diversity—matters when forecasting the impact of interventions for managing resistance.

We consider two alternative non-serotype-specific vaccines: an “acquisition-blocking” vaccine, which prevents carriage from being established with probability ε_a_, and a “clearance-accelerating” vaccine, which shortens the duration of carriage by a fraction ε_c_. Both vaccines reduce pneumococcal transmission through alternative modes of host immunity that might be elicited by a whole-cell or protein-based universal pneumococcal vaccine. Analogously to naturally-acquired serotype-independent pneumococcal immunity (*49*), the protective effect of whole-cell vaccines manifests as accelerated clearance (*50*); it is unclear whether protein-based vaccines would block pneumococcal acquisition, like PCVs, or accelerate clearance (*51*). We refer to ε_a_ or ε_c_ as the vaccine efficacy, and for simplicity, we assume that all children under five years old have vaccine protection, as would be established by an infant vaccination programme rolled out across Europe. In order to compare these vaccines with an alternative intervention of antibiotic stewardship, we also evaluate the impact of reducing the rate of penicillin prescribing by a fraction ε_s_.

We find that both vaccines have a similar impact upon carriage prevalence, regardless of whether competition or diversity maintains stability in resistance evolution (Fig. 3a). Specifically, as the vaccine efficacy ε_a_ or ε_c_ increases, carriage decreases, with the elimination of pneumococcal carriage occurring at a vaccine efficacy between 50 and 60%. Reducing antibiotic prescribing moderately increases pneumococcal carriage, such that carriage prevalence increases to approximately 54% across all countries when penicillin prescribing is eliminated completely.

**Fig. 3.**
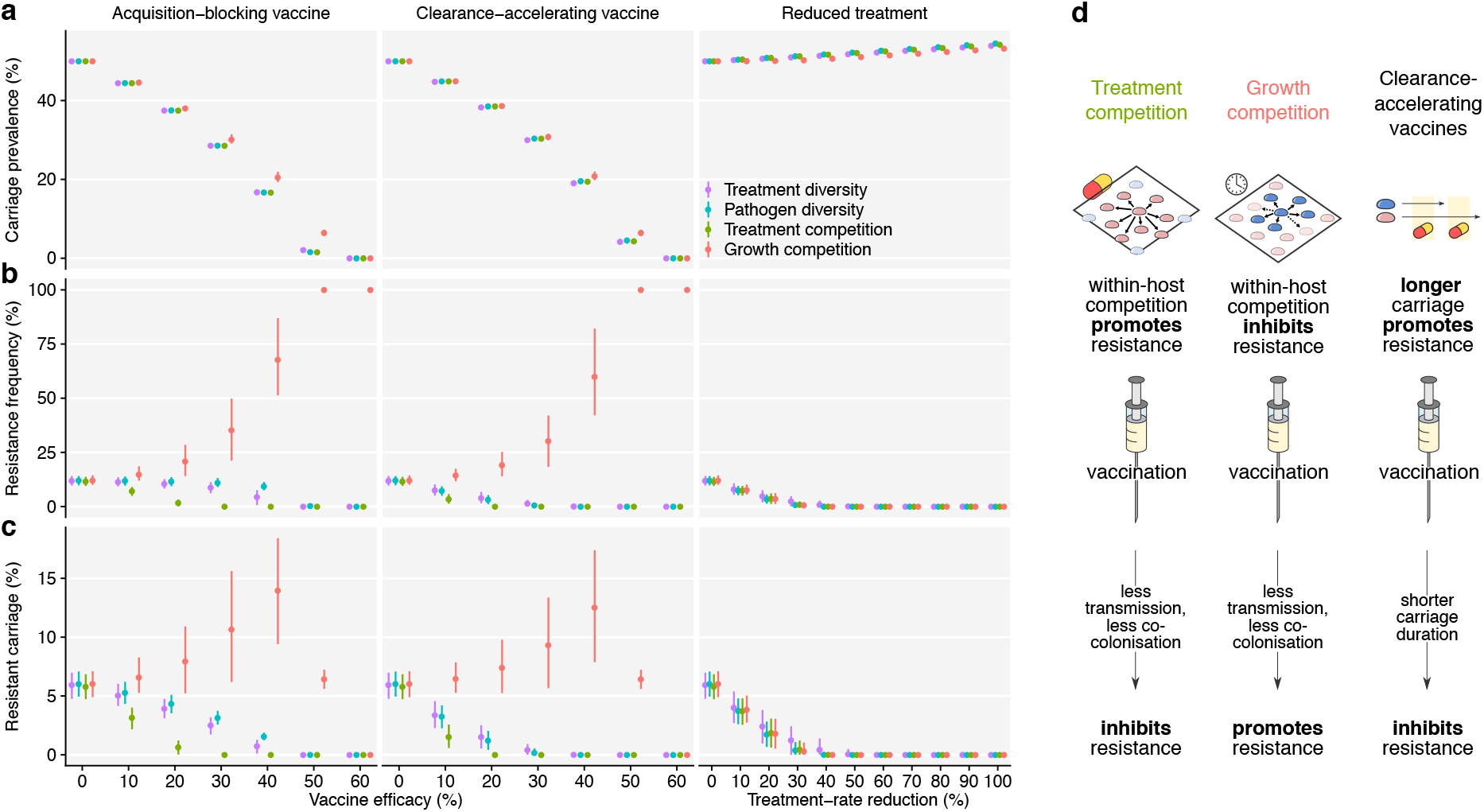
Impact of interventions. Impact of vaccine and treatment interventions on **(a)** carriage prevalence, **(b)** resistance frequency, and **(c)** resistant carriage (mean and 95% HDI). **(d)** Illustration of the strongest forces selecting for greater or lesser resistance across models.

However, the mechanism of resistance evolution has a substantial impact upon whether vaccines increase or decrease the frequency of resistance in *S. pneumoniae* in the long term (Fig. 3b). In the “Treatment diversity” and “Pathogen diversity” models, the acquisition-blocking vaccine has relatively little impact upon the frequency of resistance, because administering a universal pneumococcal vaccine to all individuals does not substantially alter the distribution of antibiotic use or of heritable variation in clearance rates. By contrast, in the within-host competition models, vaccination has a substantial impact upon resistance evolution because by reducing pneumococcal circulation, vaccines decrease the rate at which strains encounter each other within hosts, and hence strongly decrease competition between pneumococcal strains. Specifically, the acquisition-blocking vaccine selects strongly against resistance in the “Treatment competition” model: since antibiotic-mediated within-host competition benefits the resistant strain in this model, the vaccine works against this competitive advantage and therefore inhibits resistance. Conversely, in the “Growth competition” model, growth-mediated competition benefits the sensitive strain, and so by reducing competition, vaccination tends to promote resistance. These results expand upon our previous finding that the rate of co-colonisation modulates resistance evolution through its impact upon within-host competition (*26*).

The clearance-accelerating vaccine exhibits similarly divergent impacts across mechanisms of resistance evolution. However, compared with the acquisition-blocking vaccine, it also has an additional resistance-inhibiting effect across all models, because a shorter duration of carriage—whether natural or vaccine-induced—selects against resistance (*38*). This suggests that vaccines that accelerate natural clearance have a particular potential for managing resistant infections. As expected, reducing the rate of penicillin prescribing selects against resistance, exhibiting a similar impact across all four models.

The impact on resistant carriage (Fig. 3c), which combines changes in the prevalence of carriage and changes in the frequency of resistance, can be treated as a proxy for the incidence of resistant infections. Overall, under the “Growth competition” model, vaccination at intermediate efficacy is expected to increase the rate of resistant carriage, and hence the number of cases of resistant disease. In other models, vaccination always reduces resistant carriage, particularly under the “Treatment competition” model. A summary of the strongest vaccine impacts is shown in Fig. 3d.

### Evidence to inform policy and vaccine trials

For vaccines to be considered an efficient means of controlling resistant infections, they must compare favourably to existing interventions, such as reducing inappropriate antibiotic use (*52*). The UK government has recently announced an initiative to reduce antibiotic consumption by 15% by the year 2024 (*52*). Our models predict that a 15% reduction in primary-care penicillin consumption would reduce carriage of penicillin-non-susceptible pneumococci from 6% to 3%. The vaccine efficacy required to yield the same effect varies considerably depending upon the mechanism of resistance evolution (Fig 4a); for example, the required vaccine efficacy is lowest under the “Treatment competition” model (ε_a_ = 11% or ε_c_ = 7%), and highest under the “Growth competition” model (ε_a_ = 52% or ε_c_ = 50%). A full comparison of vaccine and stewardship interventions would require accounting for the economic cost of vaccines versus antibiotics, the wider range of resistant pathogens that would be targeted by restrictions on antibiotic use, and any potential increase in pathogen circulation that might be brought about by inadvertent decreases in appropriate antibiotic use.

**Fig. 4.**
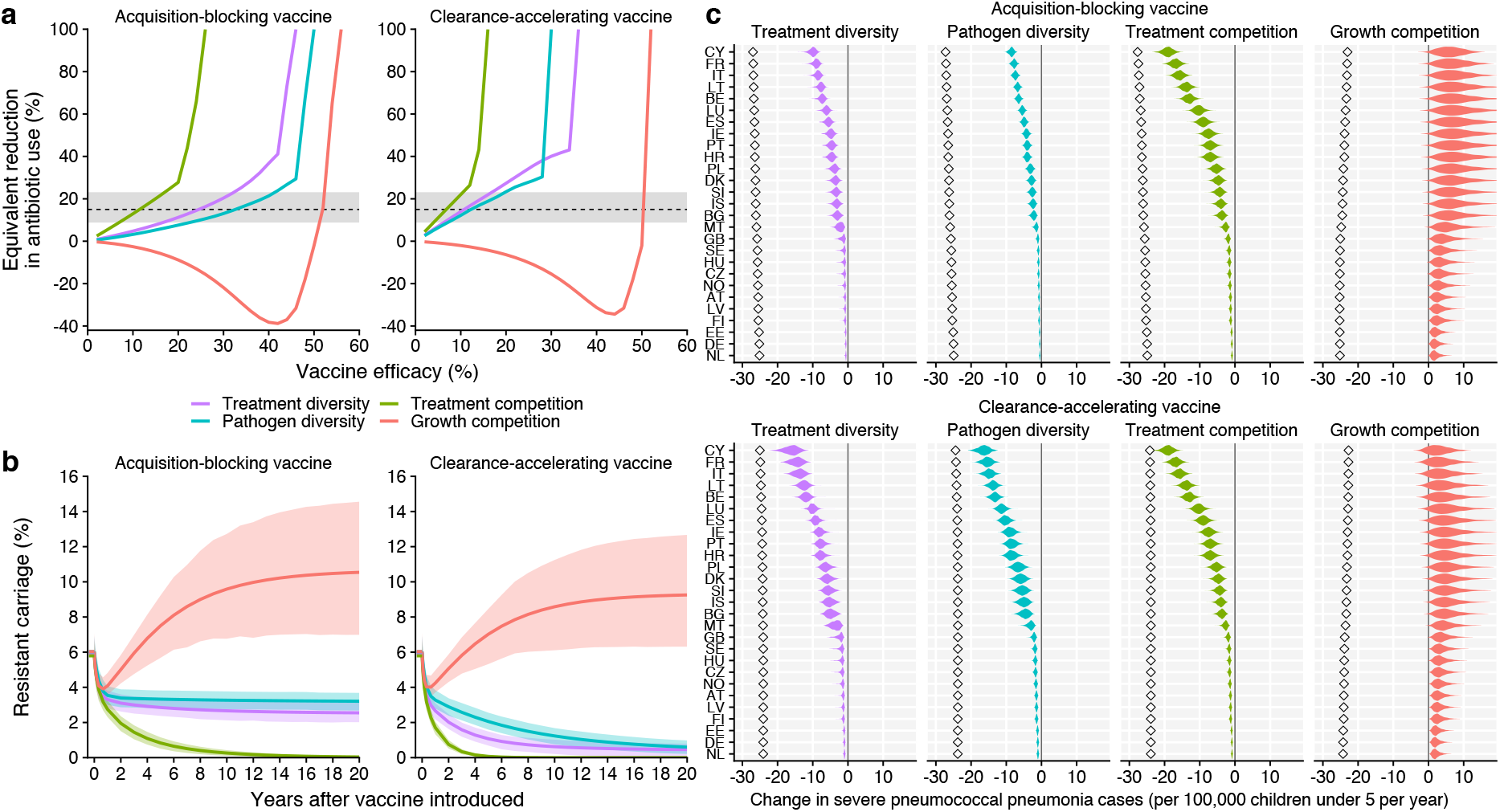
Policy considerations. **(a)** Median equivalent reduction in prescribing across four models of resistance evolution, in terms of vaccine efficacy at reducing the prevalence of resistant pneumococcal carriage. This demonstrates the vaccine efficacy required to achieve a similar decrease in resistant carriage to a given reduction in antibiotic prescription rates. The impact on overall pneumococcal carriage is not considered here. The shaded bar shows an 8.8-23.1% reduction in prescriptions, an estimate of the percentage of prescriptions which are clinically inappropriate in the UK (*77*). The dashed line shows a 15% reduction in prescriptions, which has recently been announced as a target by the UK government (*52*). **(b)** The full impact of vaccination, illustrated here with 30% vaccine efficacy, can take 5-20 years to play out (mean and 95% HDI). **(c)** Per-country impact of vaccination at 30% efficacy. Countries reporting to ECDC are ordered from lowest (NL) to highest (CY) reported rate of penicillin consumption. Diamonds show the estimated change in all pneumococcal pneumonia cases, while filled distributions show the change in resistant cases.

In randomized controlled trials of pneumococcal vaccines, resistance-related endpoints have routinely been evaluated over a follow-up period of between 6 months and 3.5 years after vaccination (*53, 54*). If vaccine-induced changes in resistance evolution unfold over a considerably longer timescale, similarly-designed trials may not fully capture vaccine impact on resistance. Indeed, we find that it can take 5-10 years for resistance to stabilise following vaccination (Fig. 4b), and that short-term drops in resistance can be reversed—or even give way to increased resistance—in the long term. Moreover, a trial in which vaccination is not offered to a substantial fraction of the population would not capture the full impact of reduced pneumococcal circulation, which is what drives competition-mediated changes in resistance in our models. Finally, our analysis assumes that vaccines are administered to all recipients simultaneously. In the real world, where vaccination is likely to be be rolled out gradually, the full effect of vaccination would take even longer to observe.

The impact of vaccination at a national level varies depending upon the treatment rate in a given country. Focusing on the specific outcome of childhood pneumococcal pneumonia cases, we find that while interventions have a consistent impact from country to country on the total pneumonia case rate, the impact on resistant pneumonia cases is greatest in those countries where antibiotic use, and hence resistance, is highest (Fig. 4c). We focus on resistant carriage, but the realised public health benefits of any intervention targeting both resistant and sensitive strains will depend upon the relative health burdens of susceptible versus non-susceptible *S. pneumoniae* infections; enumerating these comparative burdens is the subject of ongoing research (*55*).

### Vaccination in a high-burden setting

High prevalences of carriage, disease, and resistance are often observed in low-income settings, and it is desirable to know whether this could substantially alter predictions of vaccine impact. As an illustrative example, a 90% pneumococcal carriage rate, with 81% of isolates resistant to penicillin, has been observed among children under five years old in western Kenya (*56*). This may be partly attributable to a longer average duration of carriage in this setting, as a 71-day mean duration of natural pneumococcal carriage has been measured in Kilifi, eastern Kenya (*57*).

To model a similar high-burden setting, we adjust model parameters estimated from European data: increasing the mean natural carriage duration, transmission rate, and treatment rate to match observed data, and ignoring mixing with any other countries *(f* = 1), while keeping other parameters the same. We find that a comparatively greater vaccine efficacy is needed to reduce the prevalence of resistant carriage in a high-burden, high-resistance setting (Fig. 5). This is particularly true under the “Growth competition” model, because in this model resistant carriage only declines as total pneumococcal carriage declines, and it is particularly difficult to reduce overall carriage in a high-transmission setting. Simultaneously, vaccination may have a comparatively greater impact in high-burden settings because of a comparatively higher incidence of disease: for example, Kenya has been estimated to have an 8.8-fold higher incidence of severe pneumococcal pneumonia than the average in Europe (*58*).

**Fig. 5.**
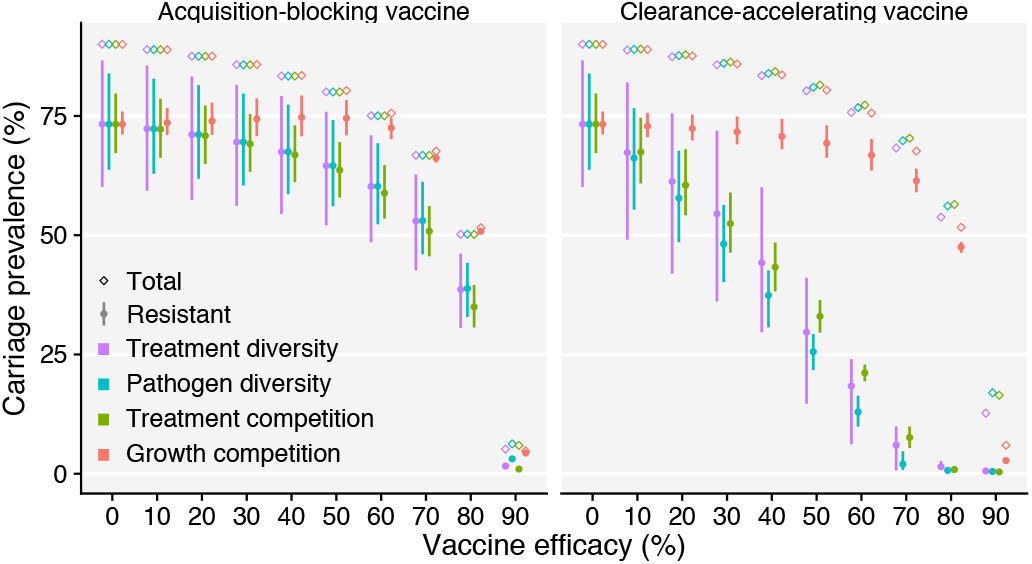
Vaccine impact in a high-burden setting. Adjusting fitted models to be consistent with a high-burden setting yields different predictions for vaccine impact (mean and 95% HDI), highlighting both increased challenges and greater opportunities for resistance management via vaccination.

### Accounting for additional between-country variation does not substantially alter predictions

Our focus thus far has been on the impact of the four identified mechanisms *per se* upon resistance evolution, and accordingly we have focused on reproducing the positive association between treatment rate and resistance frequency rather than attempting to capture the additional variability in resistance frequency between countries not accounted for by the reported treatment rate alone (Fig. 2a). This additional variability may partially stem from differences in national testing and reporting practices, or between-country differences in the distribution of pneumococcal serotypes among invasive isolates (*59*). However, another possibility is that this additional variability in resistance results from systematic differences in pathogen biology or host behaviour across countries which can be captured by our modelling framework.

To help identify which model parameters could account for this variability, we relax the assumption that only the treatment rate varies across countries, and perform Bayesian maximum *a posteriori* fitting, assuming one additional parameter *(c, b*, β, *u, f, z, g*, κ, *a*, δ, or *k*) is free to vary between countries while other parameters are held constant. We find that additional variation in resistance between countries can be explained by variation in certain other parameters, depending upon which model is used (Fig. 6a-b). Importantly, among those parameters for which additional variation between countries can explain the variation in resistance (Fig. 6c), predictions for the overall impact of vaccination remain similar, with the major differences between scenarios still attributable to the underlying mechanism of resistance evolution (Fig. 6d; Figs. S5–S16). Models that could make more accurate country-specific predictions would need to account for the effects of demographic structure, differences in carriage prevalence and disease rates between settings, and variable vaccine protection among individuals.

**Fig 6.**
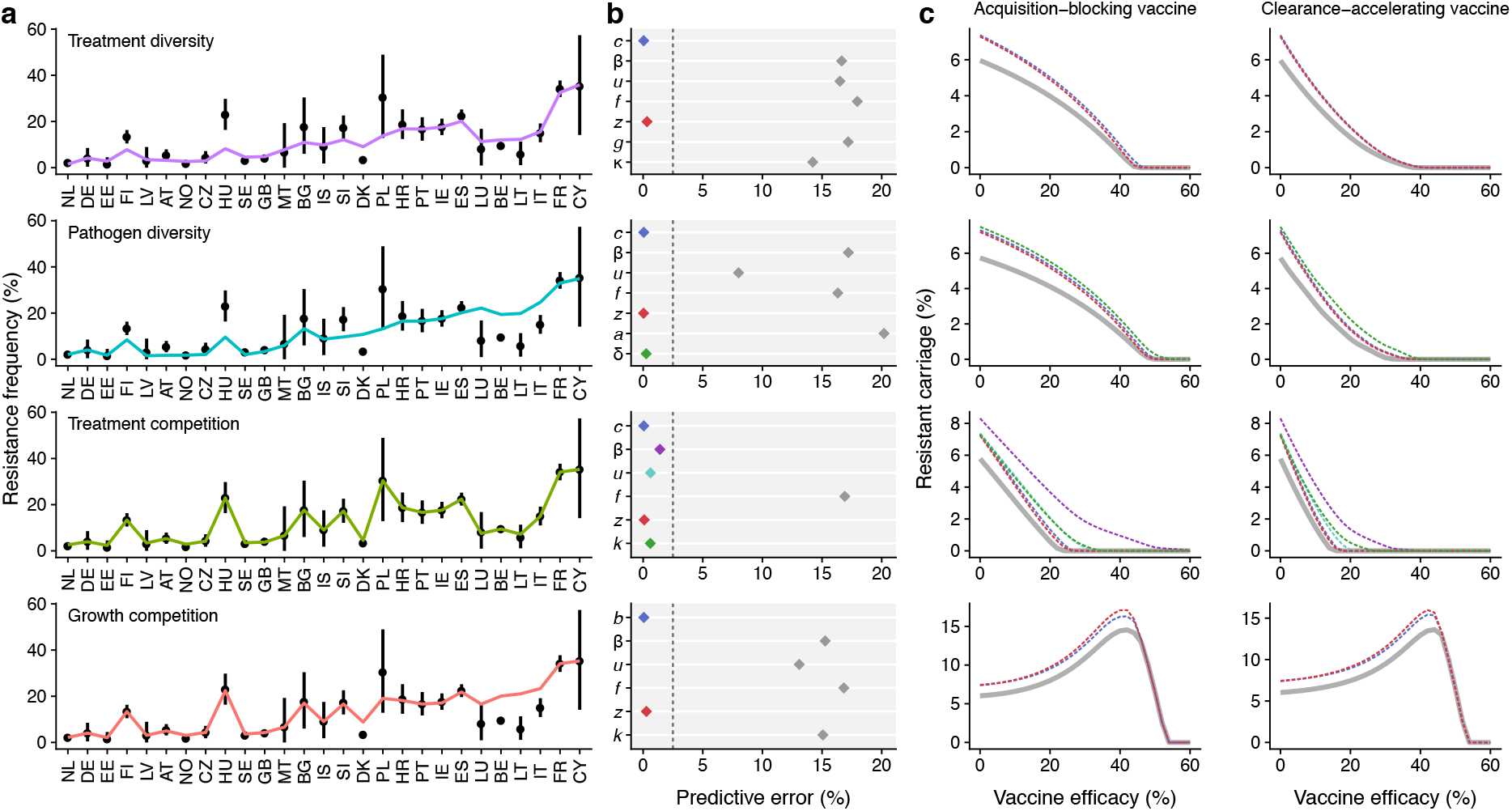
Explaining additional between-country variation in resistance frequency. Allowing model parameters to vary across countries captures additional between-country variation in resistance frequency not captured by variation in the treatment rate. For example, **(a)** allowing the transmission rate β to vary across countries can explain the variation in some but not all models (points and vertical lines show mean credible resistance frequency and 95% HDI for each country). **(b)** Depending upon which parameter is allowed to vary, models differ in how well they explain all additional between-country variation, with a clear separation (dashed line) between flexible and inflexible models. **(c)** Model-specific predictions for the impact of vaccination among those parameters that do fully capture the observed variation remain similar. Solid grey lines show “base” model; dashed lines correspond with colours in panel b.

## Discussion

We have identified four mechanisms of resistance evolution that are capable of recapitulating the observed relationship between penicillin consumption and penicillin non-susceptibility in *S. pneumoniae* across Europe. These mechanisms are not mutually exclusive, but the relative importance of each is predicted to have a substantial impact upon predictions for resistance evolution under vaccination. In particular, the “directionality” of within-host competition—that is, whether, on average, within-host competition tends to benefit resistant or sensitive strains—strongly determines whether vaccination selects for a decrease or an increase in antibiotic resistance in the long term. This directionality may vary between pathogens, but is also sensitive to the antibiotic treatment rate, and so may also vary between settings. Although we have focused on competition between sensitive and resistant strains of *S. pneumoniae* only, competition between serotypes (*24*) and with other bacteria colonizing the nasopharynx will also impact upon resistance evolution, and determining the importance of these other sources of within-host competition is crucial.

A key result of our models is that the mode of vaccine protection—whether acquisition-blocking or clearance-accelerating—has an appreciable impact upon resistance evolution. Whole-cell and purified-protein pneumococcal vaccines may induce antibody-mediated humoral immunity, CD4+ T helper-17 cell-mediated immunity, or both, with the type of immunity mediating pneumococcal acquisition, carriage, and disease in ways that are still not fully understood (*49–51*). By modelling both modes of vaccine action, we have highlighted that clearance-accelerating vaccines have increased potential for preventing the spread of resistance, because in shortening the duration of asymptomatic carriage they limit the fitness advantage of resistant pathogens under selection pressure from antibiotic use.

We fit our models to a “snapshot” of penicillin non-susceptible *S. pneumoniae* as observed across European countries in 2007, finding that each model recapitulated the data equally well. This raises the question of what kind of data would be needed to distinguish the models. One possibility would be to consider trends of resistance evolution over time. Indeed, the prevalence of penicillin non-susceptibility in *S. pneumoniae* remained largely stable in Europe between 2005-2017, a period which saw the incorporation of PCV into the routine immunization schedules of most European countries (*60*). This could be viewed as favouring the “diversity” mechanisms, which predict little change in resistance evolution following vaccination. But because serotype replacement has largely negated any vaccine impact on the prevalence of nasopharyngeal pneumococcal carriage (*10, 11*), it is not clear that we would be able to detect any effects of competition-mediated resistance evolution following a serotype-specific vaccine such as PCV—particularly given the complexity of detecting vaccine-attributable changes in resistance in a population-level associational study that would be confounded both by serotype replacement and by other changes in resistance evolution that might be expected to occur at a national level over the course of multiple years. However, it is known that the prevalence of pneumococcal carriage declines substantially with age (*42*). Therefore, it might be possible to detect a signal of within-host competition between sensitive and resistant strains by comparing the relative prevalence of resistant pneumococcal carriage in younger versus older hosts, provided that other differences could be controlled for.

Under the “Treatment diversity” and “Pathogen diversity” models, we have argued that universal pneumococcal vaccination will have little impact upon the long-term evolution of antibiotic resistance because it does not change the sources of diversity that modulate resistance evolution. Nonetheless, it is possible to target vaccines such that this diversity is harnessed to manage resistance: high-resistance serotypes could be targeted with a serotype-specific vaccine, or high-treatment subpopulations could be targeted for vaccination in order to more effectively manage resistance. Indeed, vaccination does have an additional inhibiting effect upon resistance in our models because of the latter effect. This inhibition occurs because the vaccine has a relatively greater impact upon transmission in populations where the prevalence of carriage is already low, which in our models occur in countries or subpopulations with more antibiotic consumption. Since these populations drive resistance more strongly, the vaccine’s comparatively greater impact in these populations tends to moderately inhibit resistance overall. We note that while previous work (*38*) has suggested that resistance evolution under a “Pathogen diversity” model results in a “stepped” resistance pattern in which D-types are either fully sensitive or fully resistant at equilibrium, we find that small amounts of mixing between populations can smooth out this pattern and allow intermediate rates of resistance within subtypes (Fig. S2). Finally, while we have framed “Treatment competition” and “Growth competition” as two distinct alternatives, they can instead be viewed as endpoints on a continuum, with possible models of resistance evolution for which both *c* > 0 and *b* > 0 lying between them. The impact of vaccination on resistance in such a model would depend upon the relative importance of treatment-mediated and growth-mediated competition.

This analysis has necessarily made simplifying assumptions. We have focused on prevalence (the fraction of individuals who are carriers) rather than incidence (the rate of new carriage episodes) of nasopharyngeal carriage in presenting our findings. There is evidence that pneumococcal disease progression is more likely to occur shortly after nasopharyngeal acquisition (*61*), suggesting that incidence may be more relevant than prevalence for predicting disease outcomes. Of particular note, recent modelling work has suggested that clearance-accelerating vaccines can increase rates of pneumococcal acquisition, if extended carriage is protective against new acquisition (*62*). However, it is not obvious how to compare rates of carriage acquisition across the models examined in this paper, particularly because co-colonisation is explicitly tracked in some but not all models. More work is required to clarify the links between acquisition, carriage, and disease across competing models of pneumococcal transmission. Additionally, we have assumed that antibiotic treatment rates among pneumococcal carriers remains constant after the introduction of a vaccine, even though treatment rates dropped in many settings following PCV introduction (*5, 9*). However, for a universal pneumococcal vaccine that reduces antibiotic treatment rates because it reduces carriage and thereby prevents antibiotic-treatable disease, any reduction in treatment will only occur among individuals who, because of vaccine protection, are not pneumococcal carriers, all else being equal. It might then be expected that treatment rates in carriers would remain equally high among those individuals for whom vaccine protection has failed—although physicians may be less inclined to prescribe antibiotics for respiratory tract infection more generally after the introduction of a new pneumococcal vaccine. Finally, we have focused on modelling children under 5 years old only. We would not expect incorporating age structure to lead to qualitatively different results, but age-related maturation of the immune system has been shown to be important for maintaining the circulation of pneumococcal serotypes (*49*) which we have abstracted here.

Our work helps resolve the question: What explains the persistent coexistence between resistant and sensitive strains of *S. pneumoniae? (25*) by demonstrating that multiple mechanisms are capable of explaining trends of resistance across European countries. Since there is empirical support for within-host competition between sensitive and resistant pathogen strains (*63–66*), heritable differences in the propensity for resistance within species (*38, 67)*, and within-country heterogeneity in antibiotic consumption rates (*68–70*), all of these mechanisms likely contribute to this pattern. Our results contextualize previous mathematical studies which have variously suggested that serotype-specific vaccination may increase (*24*), decrease (*22*) or have no impact upon (*18*) the frequency of resistance in *S. pneumoniae.* While the potential for vaccination to promote resistance because of competition between sensitive and resistant strains has been described previously (*24*), we have shown that vaccination can either promote or inhibit resistance depending upon the directionality of within-host competition. While vaccines targeting highly-resistant serotypes can decrease resistance (*22*), we have shown that a serotype-independent vaccine promoting accelerated natural clearance can decrease resistance across all circulating subtypes. And where single-population models have found no long-term impact of vaccination on resistance frequency (*18*), we have shown that in multi-population models, vaccination can inhibit resistance if it has a larger impact in subpopulations that consume more antibiotics. The direction and magnitude of this effect would depend upon variation in vaccine uptake, vaccine efficacy, and pathogen transmission among subpopulations, and we have not systematically explored this variation here.

A highly efficacious serotype-independent pneumococcal vaccine can indeed reduce the overall burden of antibiotic-resistant pneumococcal infections. However, the long-term effect upon resistance of a vaccine with intermediate efficacy is less certain, as vaccine impact depends crucially upon the mechanisms that drive resistance evolution. Thus, empirical investigation of pathogen competitive dynamics—and the impact of setting-specific factors on these dynamics—is needed to make accurate predictions of vaccine impact on resistant infections.

## Methods

### Study design

This study comprises four parts: a literature search used to identify plausible mechanisms through which coexistence can be maintained between sensitive and resistant pneumococcal strains across a range of antibiotic treatment rates; a mathematical modelling study embedding these mechanisms of resistance evolution in four models of pneumococcal transmission; a Bayesian statistical analysis to fit these models to empirically observed frequencies of penicillin non-susceptibility and community penicillin consumption across 27 European countries for the year 2007; and a vaccine impact analysis using these fitted models to forecast the impact of a universal pneumococcal vaccine. We use data from 2007 because changes in pneumococcal resistance reporting standards for some countries after this year hamper the between-country comparability of data (*71*). Our objectives were to identify the mechanisms potentially responsible for maintaining coexistence between resistant and sensitive pneumococci in Europe, and to determine whether the impact of vaccination on the evolution of resistance depends upon which mechanism is assumed to operate.

### Mechanisms driving resistance

We searched PubMed using the terms: (AMR OR ABR OR ((antimicrobial OR antibiotic) AND resist*)) AND ((model OR modelling OR modeling) AND (dynamic* OR transmi* OR mathematical)) AND (coexist* OR intermediate). This yielded 93 papers (Table S1). We included all papers containing a dynamic host-to-host pathogen transmission model analysing both sensitive and resistant strains with stable coexistence as an outcome of the model. From the 11 studies meeting these criteria, we identified nine unique mechanisms, two of which correspond to alternative parameterisations of a within-host competition model. We ruled out four mechanisms because of implausibility or because previous work shows that the mechanism does not bring about substantial coexistence, leaving four mechanisms (Table 1).

### Model framework

We analyse the evolution of antibiotic resistance by tracking the transmission of resistant and sensitive bacterial strains among hosts in a set of *M* countries indexed by *m* ∈ {1,2,…, *M*} using systems of ordinary differential equations.

In a simple model, hosts can either be non-carriers (X), carriers of the sensitive strain (S), or carriers of the resistant strain (R). Omitting country-specific subscripts *m* for concision, model dynamics within a country are captured by

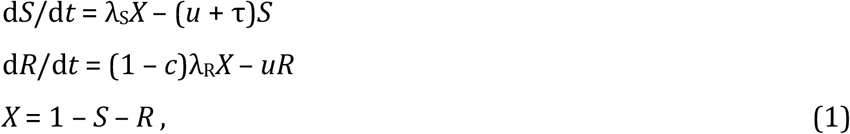

where λ_s_ is the force of infection of the sensitive strain, λ_R_ is the force of infection of the resistant strain, *c* is the transmission cost of resistance, *u* is the rate of natural clearance, and *τ* is the treatment rate. In this model, in a given country, the total carriage of the sensitive strain is *S* and the total carriage of the resistant strain is *R*. Force of infection terms are defined below; a summary of all model parameters can be found in Table S2.

The “Treatment diversity” model extends the simple model (eq. 1) by structuring each country into multiple subpopulations that exhibit different rates of antibiotic treatment and make contact with each other at unequal rates (*25, 34, 35, 72*). In each country, we model *N* equally-sized representative subpopulations indexed by *i* ∈ {1,2,…, *N*}, where we assume *N* = 10. Dynamics within a country are

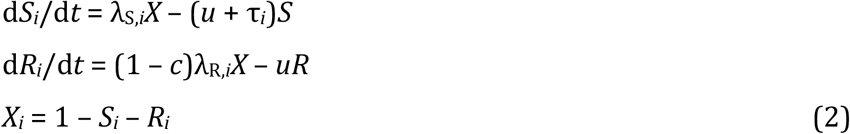

where we assume that treatment rates of subpopulations within a country approximately follow a gamma distribution with shape parameter *κ* and mean treatment rate τ. Accordingly, the rate of antibiotic consumption in subpopulation *i* is 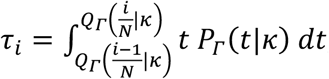, where *Q*_Γ_ *(q* |κ) is the quantile *q* of the gamma distribution with shape *τ* and *P*_Γ_ (*t* | *τ*) is the probability density at *t* of the same gamma distribution. At the scale of individuals, antibiotic consumption is highly variable, with some people taking no antibiotics in a given year and others taking many courses of antibiotics (*73*); at regional scales, antibiotic consumption shows less extreme variability (*74*) and approaches a normal distribution. We use a gamma distribution to model variation in treatment rates among subpopulations because it can capture patterns at either of these scales, or scales in between.

The “Pathogen diversity” model extends the simple model (eq. 1) by structuring the pathogen population into *D* different “D-types” (we assume *D* = 25), each with a different natural clearance rate, where each type is kept circulating by diversifying selection acting on D-type (*38*). Dynamics within a country are

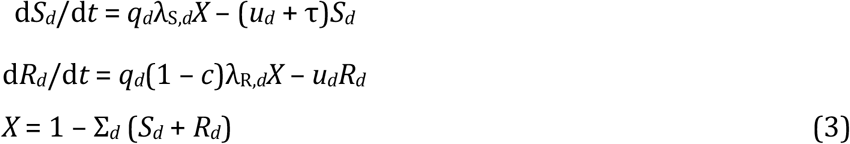

where *q_d_* is the strength of diversifying selection for D-type *d* ∈ {1,2,…, *D*} and *u_d_* is the clearance rate for D-type *d*. We follow Lehtinen *et al.* (38) in defining 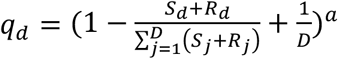 and 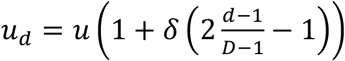, where *a* is the power of diversifying selection and *δ* is the range of clearance rates. In a given country, the total carriage of the type-*d* sensitive strain is *Sd* and the total carriage of the type-*d* resistant strain is *R_d_*.

Finally, the within-host competition models (*26*) allow hosts to carry a mix of both strains. Hosts can carry the sensitive strain with a small complement of the resistant strain (S_R_) or the resistant strain with a small complement of the sensitive strain (R_S_). Dynamics within a country are

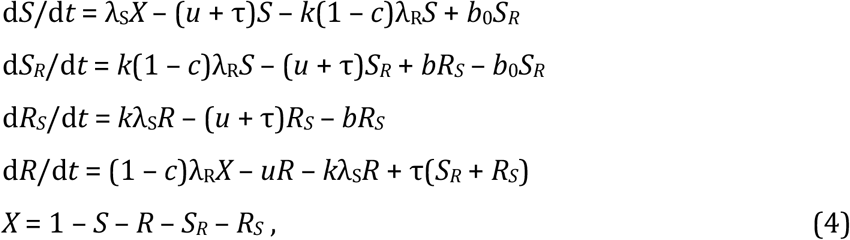

where *k* is the rate of co-colonisation relative to primary colonisation, *b* is the within-host growth benefit of sensitivity (i.e. the rate of the R_S_ → S_R_ transition), and *b*_0_ is the rate of the S_R_ → S transition. We follow Davies *et al. (26*) in setting *b*_0_ = 4*b*. In a given country, the total carriage of the sensitive strain is *S* + *S_R_* and the total carriage of the resistant strain is *R* + *R_S_*. “Treatment competition” assumes the cost of resistance is incurred by reduced transmission potential (*b* = 0 and *c* > 0), while “Growth competition” assumes that the cost of resistance is incurred through decreased within-host growth (*b* > 0 and *c* = 0).

In equations 1, 3 and 4, the force of infection of a particular strain *A* in country *m* is 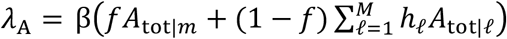, where β is the transmission rate, *f* is the between-country assortativity, *h*_ℓ_ is the relative population size of country *m* (such that *∑_ℓ_ h_ℓ_* = 1), and *A_tot|ℓ_* is the total carriage of strain *A* in country *ℓ*. The probability with which individuals contact an individual from another country, 1 - *f*, captures those contacts made with individuals from another country in either one’s home country or a foreign country. In equation 2, the force of infection of a particular strain *A* in subpopulation *i* of country *m* is 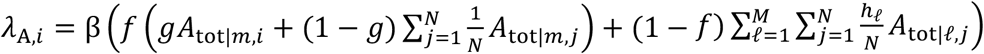, where *g* is the within-country assortativity and A_tot|*ℓ,j*_ is the total carriage of strain *A* in subpopulation *j* of country *ℓ*.

### Data and model fitting

We extracted community penicillin consumption and penicillin non-susceptibility in *S. pneumoniae* invasive isolates from databases made available by the ECDC (*13, 28*). We assume that community penicillin consumption drives penicillin resistance, that antibiotic consumption is independent of whether an individual is colonised by pneumococcus, and that resistance among invasive bacterial isolates is representative of resistance among circulating strains more broadly. Countries report community penicillin consumption in defined daily doses (DDD) per thousand individuals per day. To transform this bulk consumption rate into the rate at which individuals undertake a course of antibiotic therapy, we analysed prescribing data from eight European countries, estimating that, on average, 5 DDD in the population at large correspond to one treatment course for a child under 5 years of age. This conversion rate varies between countries (Table S3), but since the data are incomplete (8 of 27 countries) we have not explicitly accounted for this variability in our main model fitting results.

Our model framework tracks carriage of *S. pneumoniae* among children aged 0-5 years, the age group driving both transmission and disease. In European countries, we assume that the prevalence of pneumococcal carriage in under-5s is 50% (*11, 42*) and the average duration of carriage is 47 days (*47, 48*). We calculate the average incidence of *S. pneumoniae-caused* severe pneumonia requiring hospitalisation as 610 per million children under 5 per year (*58*) across the European countries in our data set. See Tables S4, S5, and S6 for details of calculations relating to pneumococcal carriage duration and disease incidence.

We use Bayesian inference via differential evolution Markov chain Monte Carlo (*75*) to identify model parameters that are consistent with empirical data while accounting for uncertainty in those estimates. Country *m* has antibiotic treatment rate *τ_m_* and reports *r_m_* of *n_m_* isolates are resistant. Over all *M* countries, these data are denoted *τ = (τ_1_, τ_2_,…, τ_M_), r* = (*r*_1_, *r*_2_,…, *r*_M_), and *n* = (*n*_1_, *n*_2_,…, *n_M_*), respectively. The probability of a given set of model parameters *θ* is then

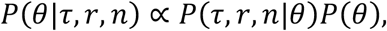

where *P*(*θ*) is the prior probability of parameters *θ* and

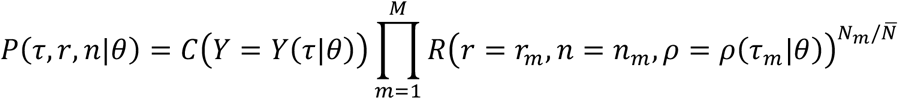

is the likelihood of data *τ, r, n* given model parameters *θ*. Above, *Y*(*θ*) is the average model-predicted prevalence of carriage across all countries and *p(τ_m_*|*θ*) is the model-predicted resistance prevalence for country *m*.*C*(*Y*) is the credibility of prevalence of carriage *Y* and *R*(*r,n,ρ*) is the credibility of *r* out of *n* isolates being resistant when the model-predicted resistance prevalence is *ρ*. For *C*(*Y*), we use a normal distribution with mean 0.5 and standard deviation 0.002. For *R*(*r*,*n*,*ρ*), we use 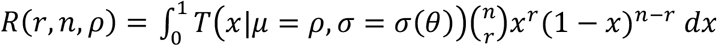 a binomial distribution where the probability of success is modelled as a [0,1]-truncated normal distribution centred on p and with standard deviation σ. The parameter σ captures the unexplained between-country variation in resistance frequency. Here, 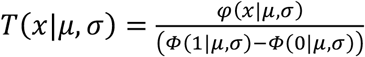, where 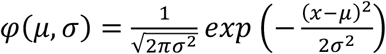 is the untruncated normal PDF and 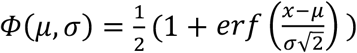 is the untruncated normal cumulative distribution function. Finally, *N_m_* is the population size of country *m* and 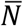 is the average population size across all countries; the exponent 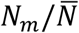 allows us to weight the importance of each country by its population size, which allows a closer fit with the overall resistance prevalence across all countries.

As prior distributions for parameter inference, we adopt *c* ~ Beta(α = 1.5, *β =* 8.5), *b ~* Gamma(κ = 2,0 = 0.5), *β ~* Gamma(κ = 5,0 = 0.35), *g ~* Beta(α = *10, β* = 1.5), *κ* ~ Gamma(κ = 4,0 = 2), *a* ~ Gamma(κ = 2,0 = 5), *δ* ~ Beta(α = 20, *β* = 25), and *k* ~ Normal(μ = 1, *σ* = 0.5). Priors for *c, b*, and β were chosen to be vague since these parameters are heavily constrained by the data we fit our models to. Priors for *g*, *κ*, *a*, δ, and *k* were chosen to keep parameters within biologically plausible ranges. Table S7 provides more detail on the choice of priors, and Fig. S4 shows which parameters are most strongly constrained by these prior beliefs.

We set the unexplained between-country variation in resistance prevalence σ to 0.06 across all models based on a preliminary round of model fitting with σ as a free parameter. We set the between-country assortativity *f* to 0.985 (*i.e.*, 1.5% of contacts occur with individuals from a different country) based on rates of travel for EU residents. Specifically, using Eurostat database tour_dem_tnw (*76*) we estimated that the average EU resident spent 1.5% of their nights abroad in 2007; this overestimates mixing because children under 5 travel less than the average person, but underestimates mixing because it does not account for contacts made with visitors to an individual’s country of residence and because children may contract pneumococcal carriage from adults who travel, and so we kept the value of 1.5%. See Table S8 for MCMC diagnostics.

To match model predictions to a high-burden setting, we increase the duration of carriage to 71.4 days; increase the transmission rate by a factor of 3.49 (Treatment diversity), 3.62 (Pathogen diversity), 3.61 (Treatment competition), or 3.20 (Growth competition), so that carriage prevalence reaches 90.0%; and increase the antibiotic consumption rate to 1.670, 1.458, 1.138, or 5.887 courses per person per year, respectively, so that resistance prevalence reaches 81.4%.

### Interventions

Interventions have the following impact on model parameters: for the acquisition-blocking vaccine, the transmission rate becomes β’ = (1-ε_a_)β; for the clearance-accelerating vaccine, the clearance rate becomes *u’* = *u*/(1-ε_c_); and under antibiotic stewardship, the average treatment rate in each country *m* becomes *τ_m_*’ = *τ_m_* (1-*ε_s_*).

### Capturing additional between-country variation in resistance frequency

We begin by finding the maximum *a posteriori* model fits according to the likelihood and prior distributions for each of the four models of resistance evolution. This identifies the following parameter values for each model. “Treatment diversity”: β = 1.41, *c* = 0.124, *g* = 0.976, and *κ* = 2.22. “Pathogen diversity”: β = 1.33, *c* = 0.191, *a* = 10.8, and δ = 0.608. “Treatment competition”: β = 1.42, *c* = 0.191, and *k* = 1.64. “Growth competition”: β = 1.39, *b* = 0.195, and *k* = 1.61. Then, we perform maximum *a posteriori* model fits for each potentially-varying parameter under each model, allowing the varying parameter to take on a different value for each country and fixing other parameters at their maximum *a posteriori* values as determined in the previous step, or at specific assumed values for *u* = 0.65, *f* = 0.985, and *z* = 5. For the second step, we use a modified likelihood function

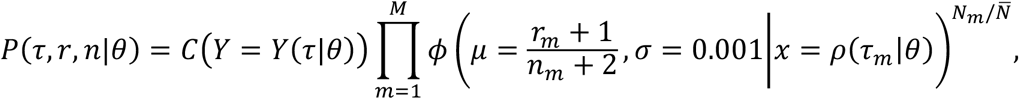

where ϕ(μ, σ|%) is the normal probability density function. This modified likelihood function ensures that the model-predicted resistance frequency for each country is matched as closely as possible to the maximum-likelihood resistance prevalence 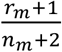 (*i.e.*, assuming a uniform prior on resistance frequency) for each country *m*, so that model fits are comparable across different varying parameters. We use the Nelder-Mead algorithm to maximize the posterior probability in both steps.

Figs. S5–S8 show maximum *a posteriori* fits when allowing an additional parameter to vary freely between countries, along with the parameter values identified by model fitting. Figs. S9–S12 show the impact of vaccination, focusing on those parameters for which model fitting was able to capture the observed variability in resistance frequency between countries (*i.e.*, those parameters plotted to the left of the dashed line in Fig. 6b of the main text). Figs. S13–S16 show the impact of vaccination for the remaining parameters.

## Author contributions

All authors designed the study. N.G.D. led the analysis with input from K.E.A., M.J., and S.F. The paper was written by N.G.D. and K.E.A. with input from M.J. and S.F.

## Code availability

C++ code used for model comparison is available at https://github.com/nicholasdavies/amr-competition-diversity.

## List of Supplementary Materials

**Fig. S1.**
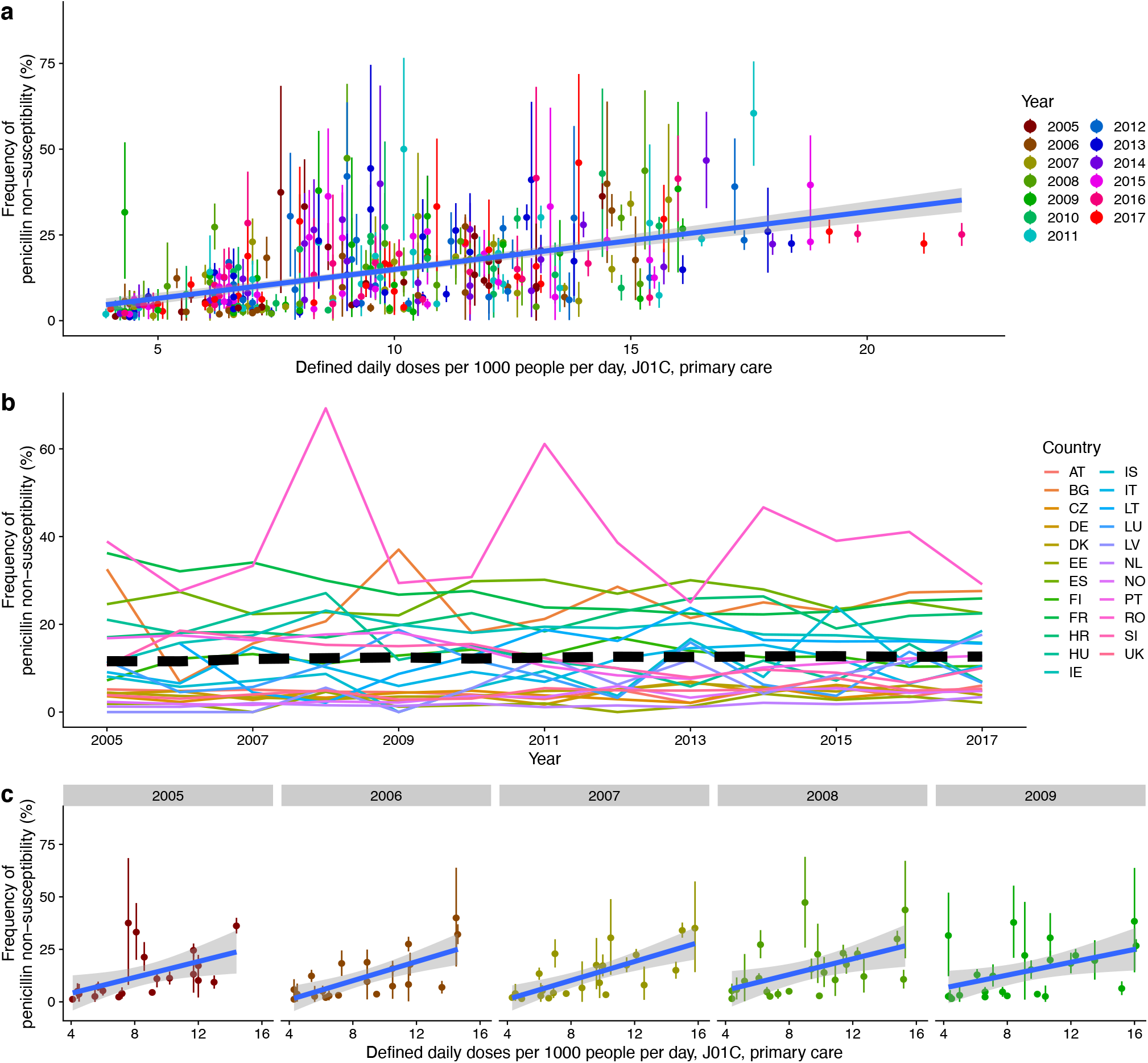
Patterns of penicillin non-susceptibility across European countries, 2005-2017. **(a)** The frequency of penicillin non-susceptibility in pneumococcal isolates (mean and 95% HDI) increases with the primary care consumption of penicillins, ATC class J01C (least-squares linear regression *β =* 0.0168, F(1,344) = 148.5, *P < 2.2 ×* 10^−16^). Data from Belgium after 2007 are excluded because of changes to the definition of penicillin non-susceptibility after this year. **(b)** The frequency of penicillin nonsusceptibility has fluctuated in individual countries between 2005 and 2017, but the European population-weighted average (thick dashed line) has remained stable at roughly 12%. **(c)** We fit our models to the prevalence of penicillin non-susceptibility in 2007. Despite the introduction of pneumococcal conjugate vaccines into many European national immunisation programmes starting in 2006, pneumococcal resistance appears to have been relatively stable over the time period 2005-2009: while mean penicillin non-susceptibility across European countries increased by 0.5% per year over this time period, and the slope of penicillin non-susceptibility on primary-care penicillin consumption decreased by 0.1%/DDD per year, neither of these changes are significant (least-squares linear regression: *F*(1,3) = 0.64, *P* = 0.48 and *F*(1,3) = 1.48, P = 0.31, respectively).

**Fig. S2.**
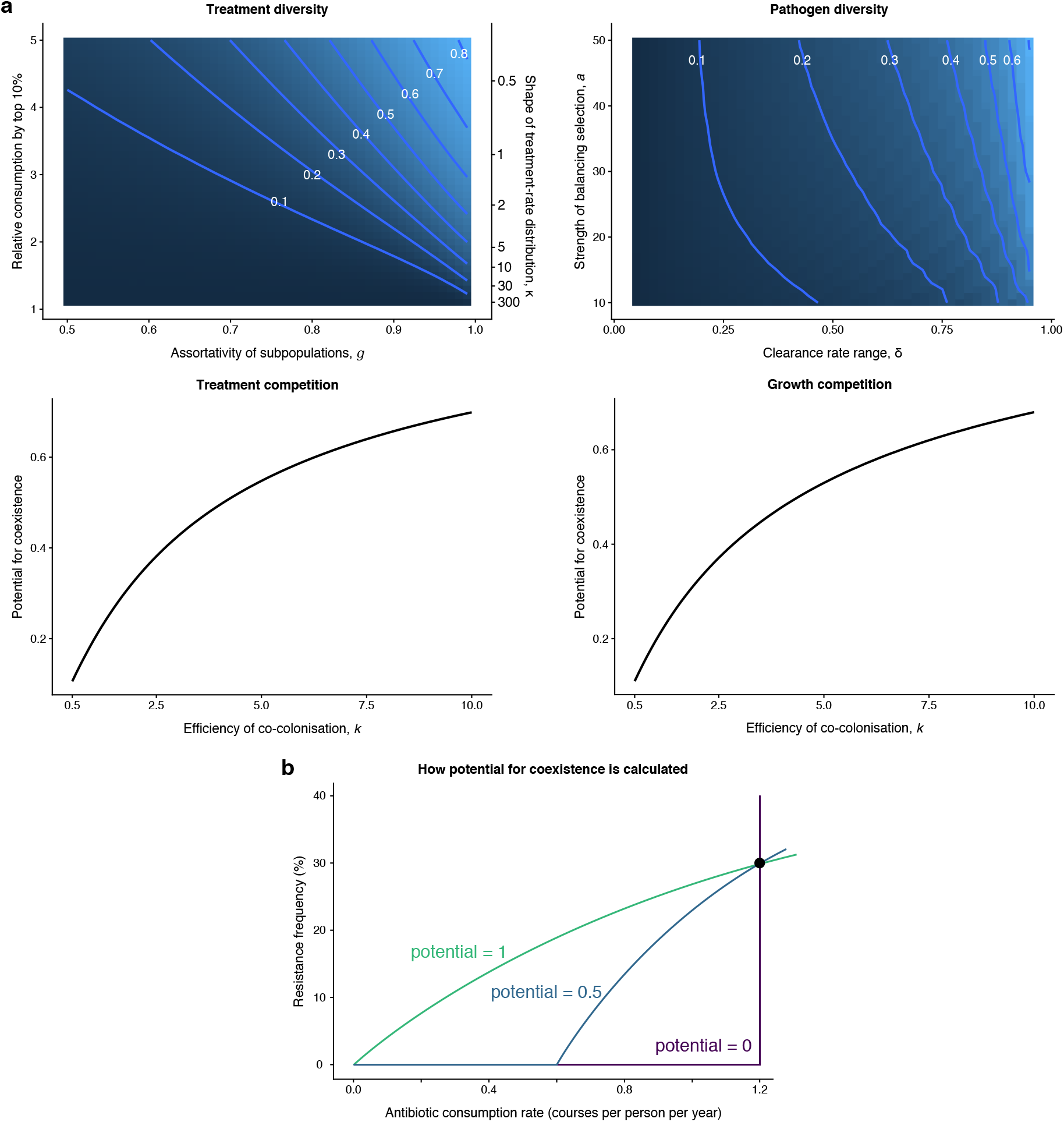
Impact of key parameters upon the potential for coexistence in each model. **(a)** Potential for coexistence for each model, depending upon key parameters. In the top two panels, darker colours represent lower potential for coexistence while lighter colours represent higher potential for coexistence. **(b)** Schematic showing how to calculate the “potential for coexistence”, an ad-hoc measurement of how much coexistence is exhibited by a given model under a given parameterisation. To find it, we set β = 1.4 months^−1^, *u* = 0.7 months^−1^, and set *g*, *κ*, *δ*, *a*, and *k* according to the values shown in the figure (panel a). We then numerically identify the value of *c* (or *b* for the “growth competition” model) which results in the equilibrium resistance frequency passing through exactly 30% for a single country in which the antibiotic consumption rate is equal to τ = 1.2 courses per person per year (black dot; these values are chosen to approximately match observations for *S. pneumoniae* in European countries). The potential for coexistence is the probability that the equilibrium resistance frequency is above 0 for a random treatment rate between τ = 0 and τ = 1.2 (in other words, it is the proportion of the curve that “lifts off” from the x-axis; see figure). By this definition, a model showing no coexistence has potential 0, while a model showing the maximum amount of coexistence has potential 1.

**Fig. S3.**
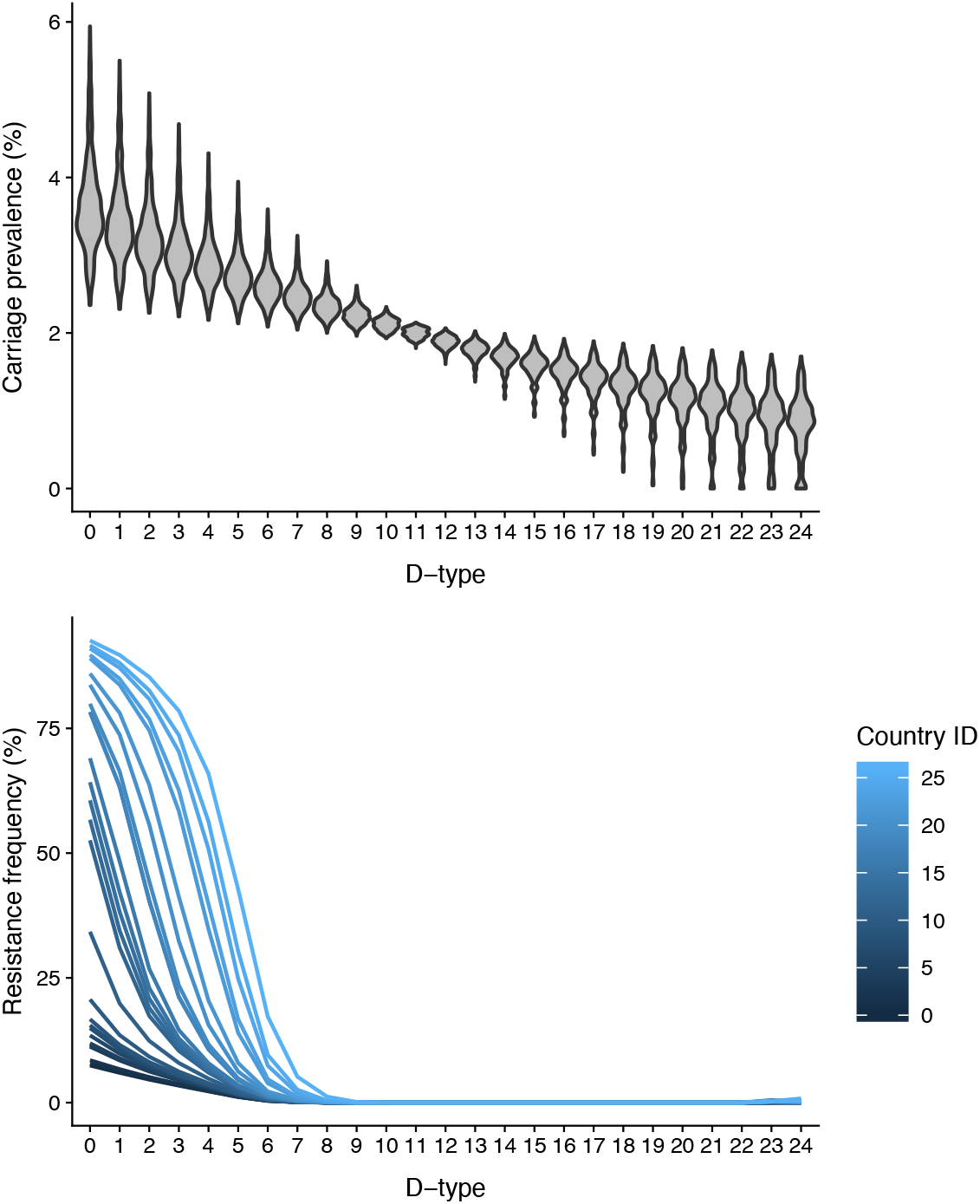
Carriage and resistance of D-types in the “Pathogen diversity” model. This verifies that the D-types with the highest prevalence of carriage (averaged over all countries, above) also exhibit the highest resistance frequency (separated by country, below), and shows that at equilibrium, D-types can exhibit intermediate frequencies of resistance.

**Fig. S4.**
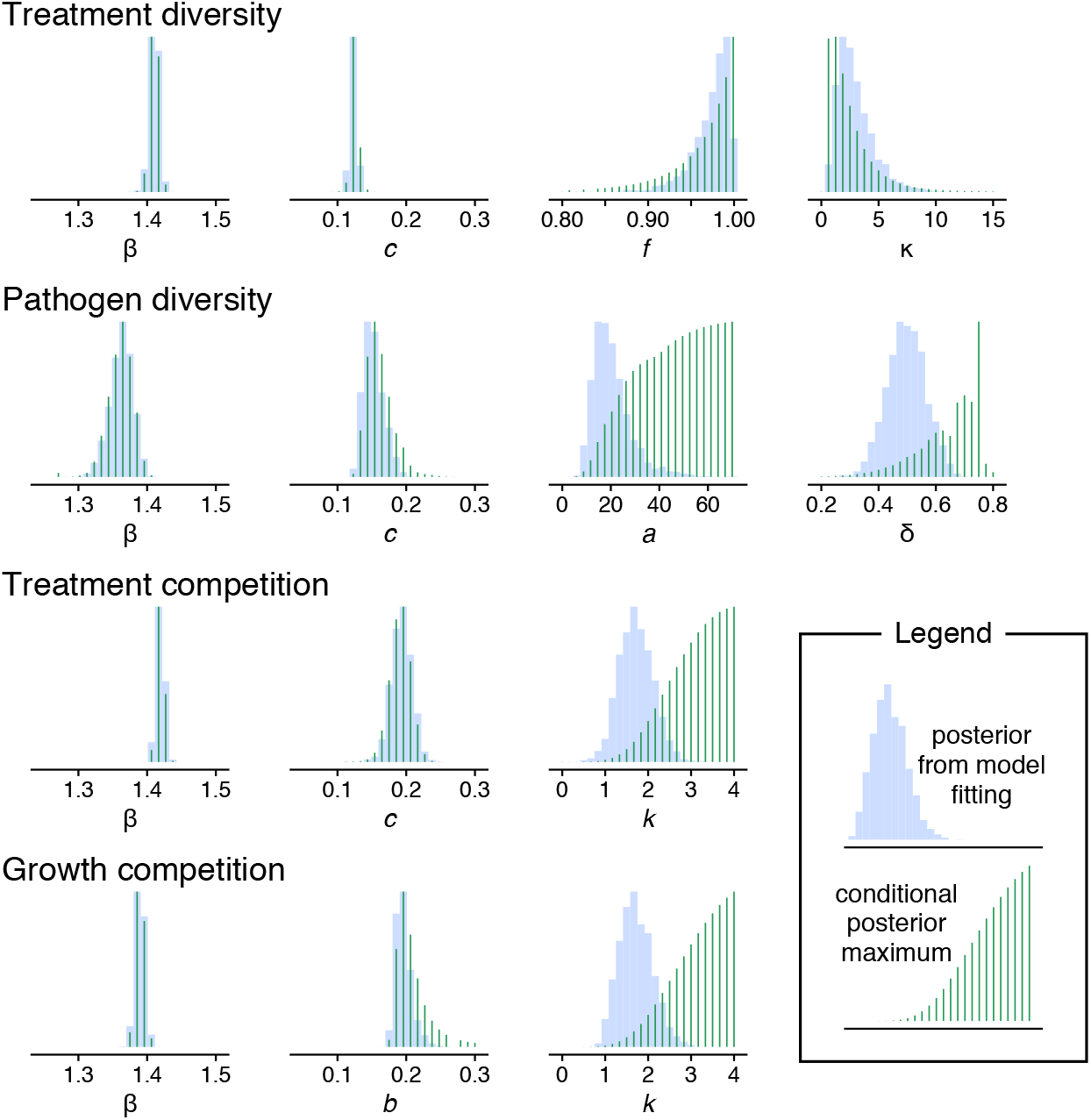
Comparison of inferred model posteriors and conditional posterior maxima for each parameter. The conditional posterior maximum (green bars) is obtained by fixing the parameter of interest at a specific value, then allowing the other parameters to assume their maximum *a posteriori* values through numerical optimization of the model fit. This is repeated for multiple values of the parameter of interest. The position on the x-axis of the thin green bars shows these fixed values for the parameter of interest (chosen to “sweep” across the parameter range), while the height of the thin green bars is relative to the maximum posterior probability conditional on that fixed value. When the conditional posterior maximum (green bars) does not align with the posteriors from model fitting (blue bars, same as Fig. 2b, main text), this indicates that the prior distribution is strongly influencing the posterior distribution for a given parameter. This analysis shows that values inferred for *a* and δ under the “Pathogen diversity” model, and for *k* under the “Treatment competition” and “Growth competition” models, are strongly constrained by prior beliefs about plausible ranges for those parameters, while other parameters are less constrained.

**Fig. S5.**
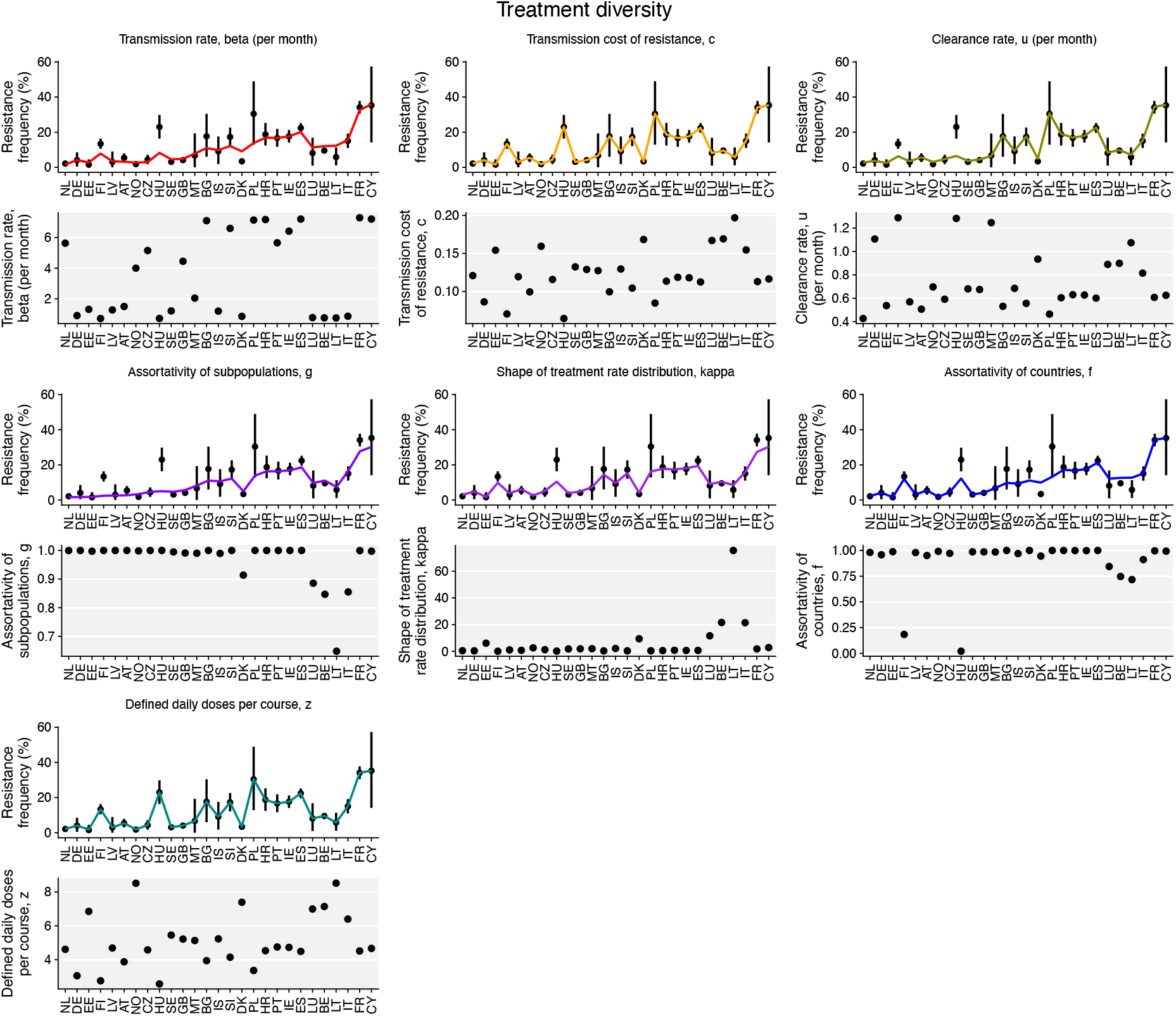
Varying-parameter fits for the “Treatment diversity” model. Maximum *a posteriori* fits for the “Treatment diversity” model allowing one parameter to vary between countries. Parameters *c* and *z* can capture the additional variation in resistance

**Fig. S6.**
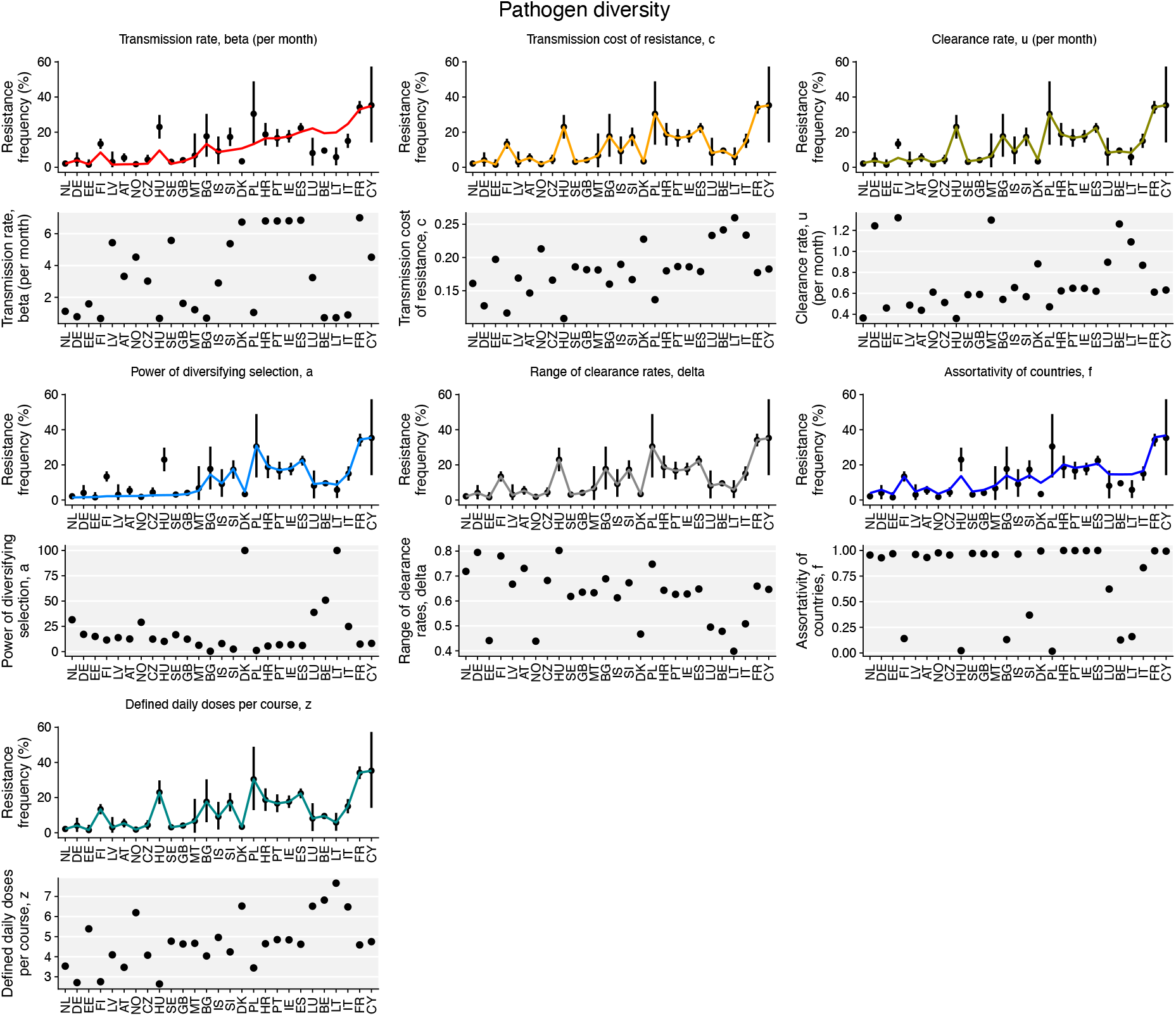
Varying-parameter fits for the “Pathogen diversity” model. Maximum *a posteriori* fits for the “Pathogen diversity” model allowing one parameter to vary between countries. Parameters *c*, δ, and *z* can capture

**Fig. S7.**
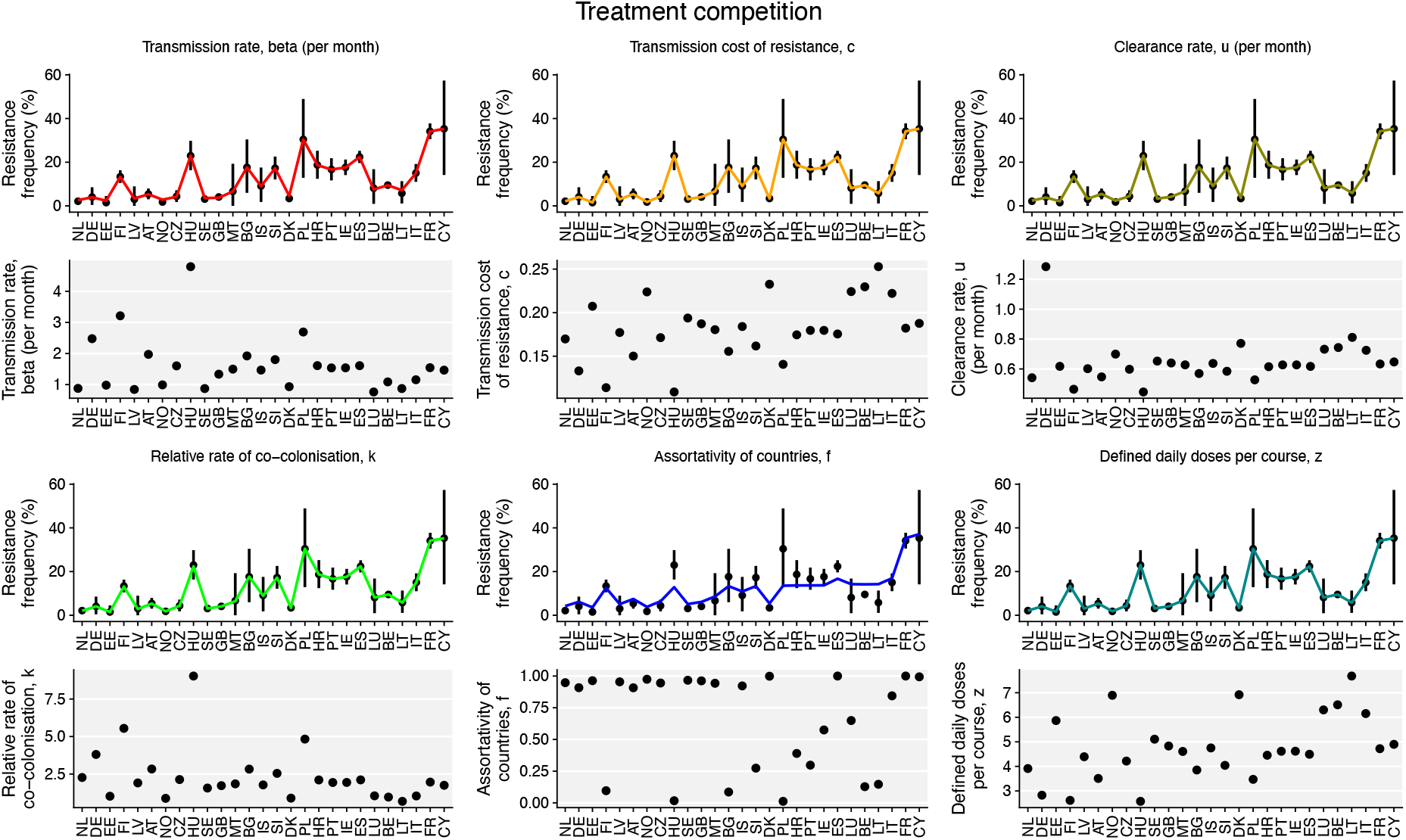
Varying-parameter fits for the “Treatment competition” model. Maximum *a posteriori* fits for the “Treatment competition” model allowing one parameter to vary between countries. Parameters β, *c*, *u*, *k*, and *z* can capture the additional variation in resistance frequency between countries.

**Fig. S8.**
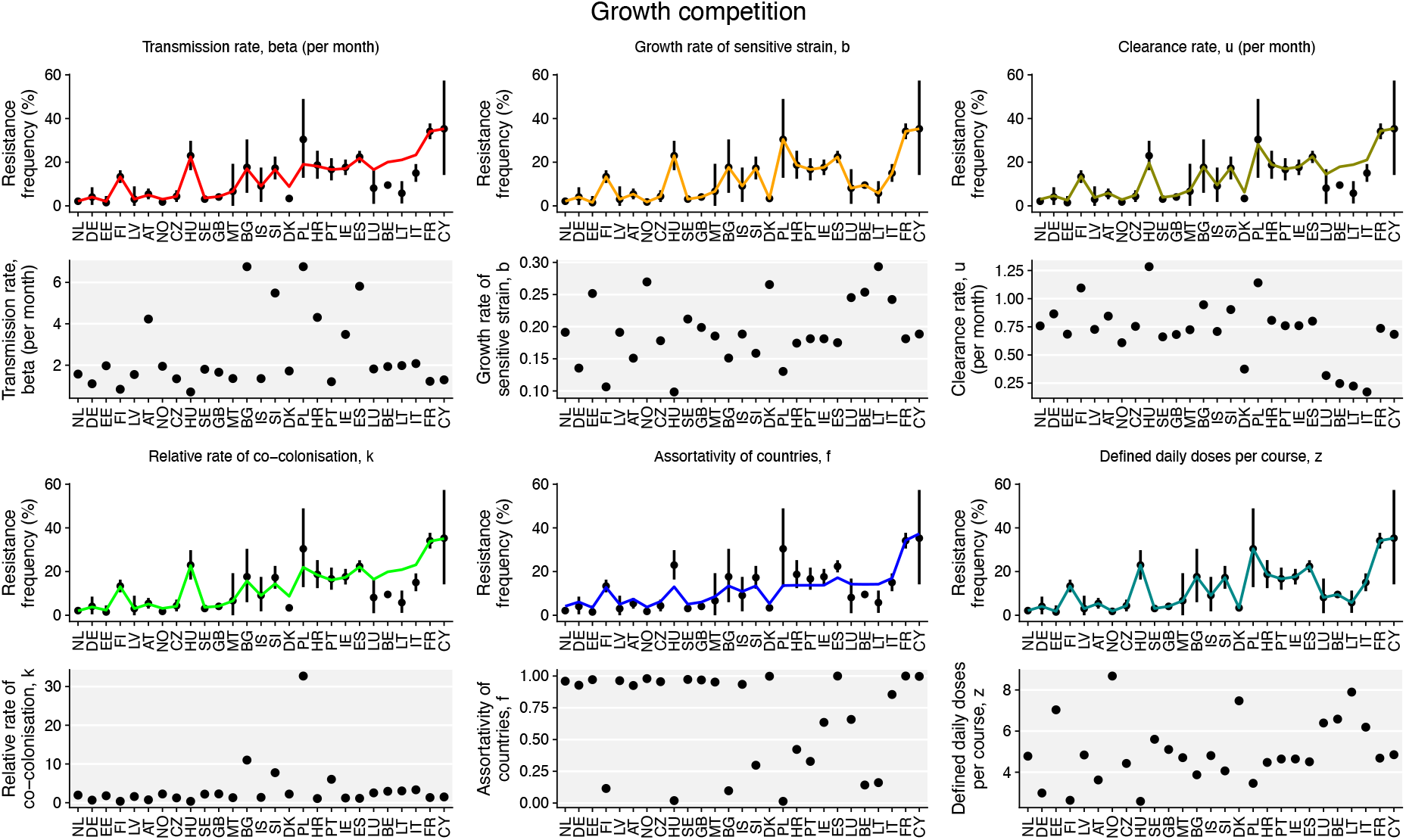
Varying-parameter fits for the “Growth competition” model. Maximum *a posteriori* fits for the “Growth competition” model allowing one parameter to vary between countries. Parameters *b* and *z* can capture the additional variation in resistance frequency between countries.

**Fig. S9.**
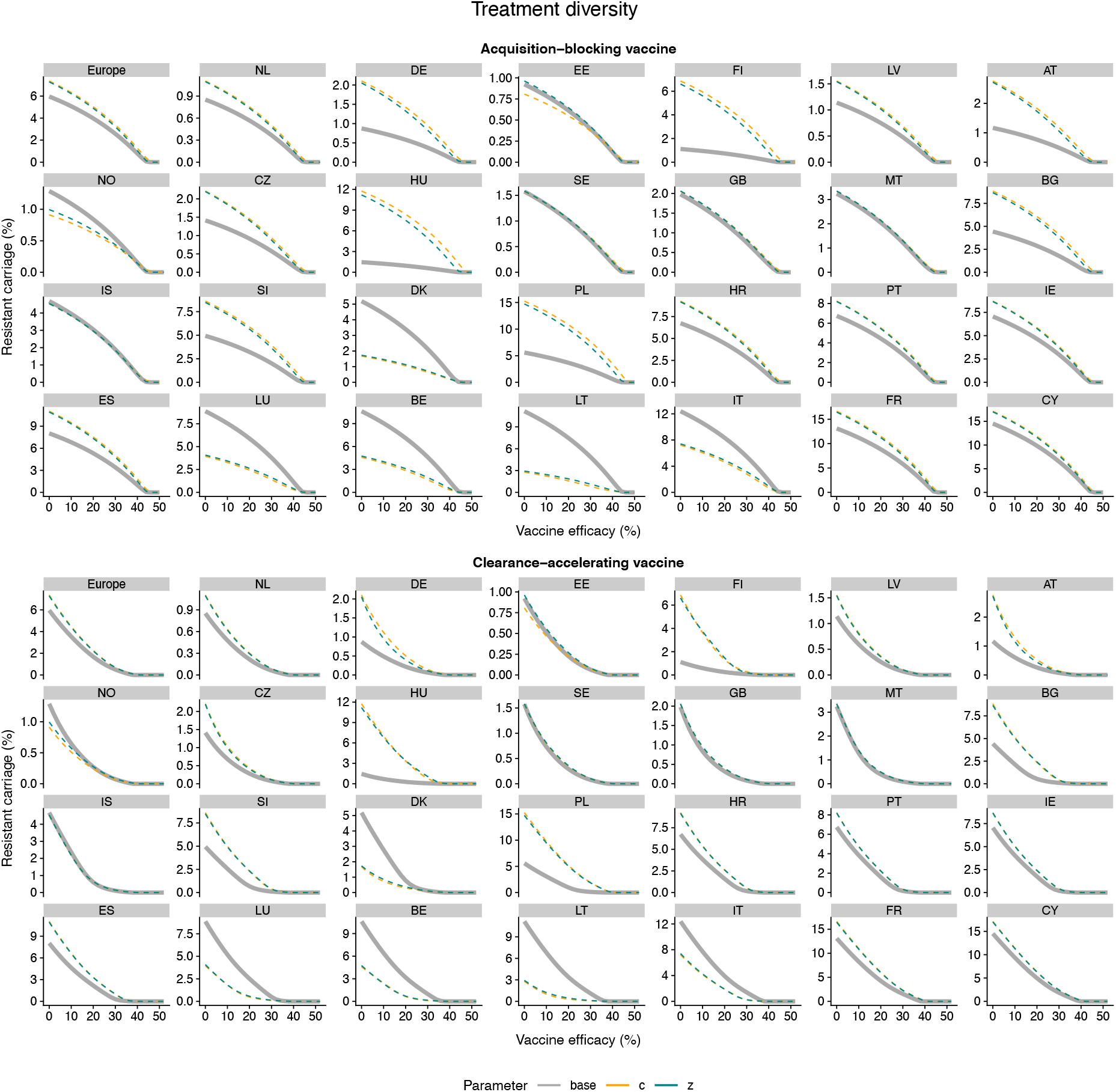
Vaccine impact for the “Treatment diversity” model, varying parameters *c* and *z.* Impact of vaccination under the “Treatment diversity” model, for those parameters able to capture the between-country variation in resistance frequency. The base model fit (thick grey solid line) is compared with the model fits in which parameters vary between countries (thin dashed lines).

**Fig. S10.**
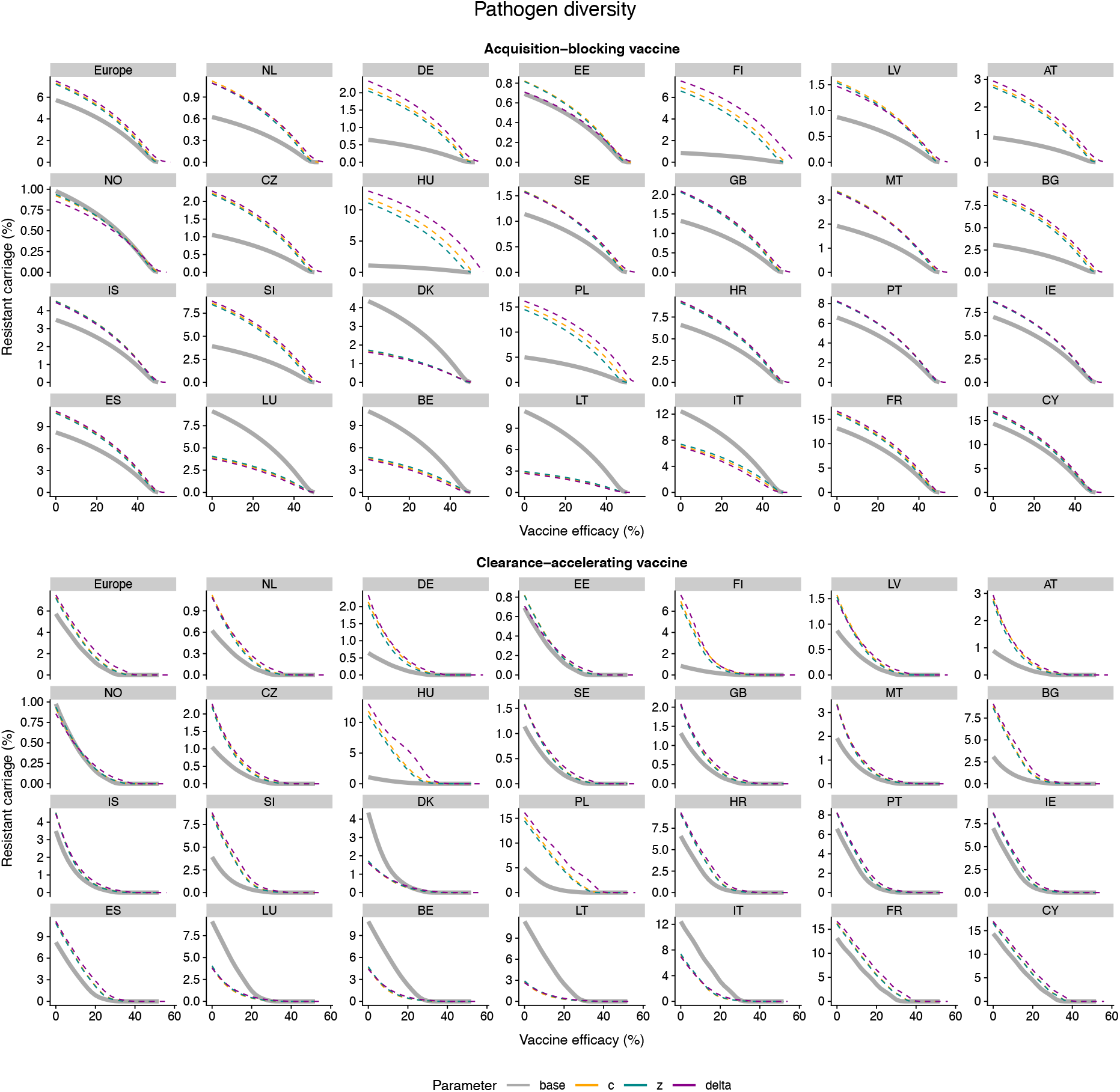
Vaccine impact for the “Pathogen diversity” model, varying parameters *c*, δ, and *z.* Impact of vaccination under the “Pathogen diversity” model, for those parameters able to capture the between-country variation in resistance frequency. The base model fit (thick grey solid line) is compared with the model fits in which parameters vary between countries (thin dashed lines).

**Fig. S11.**
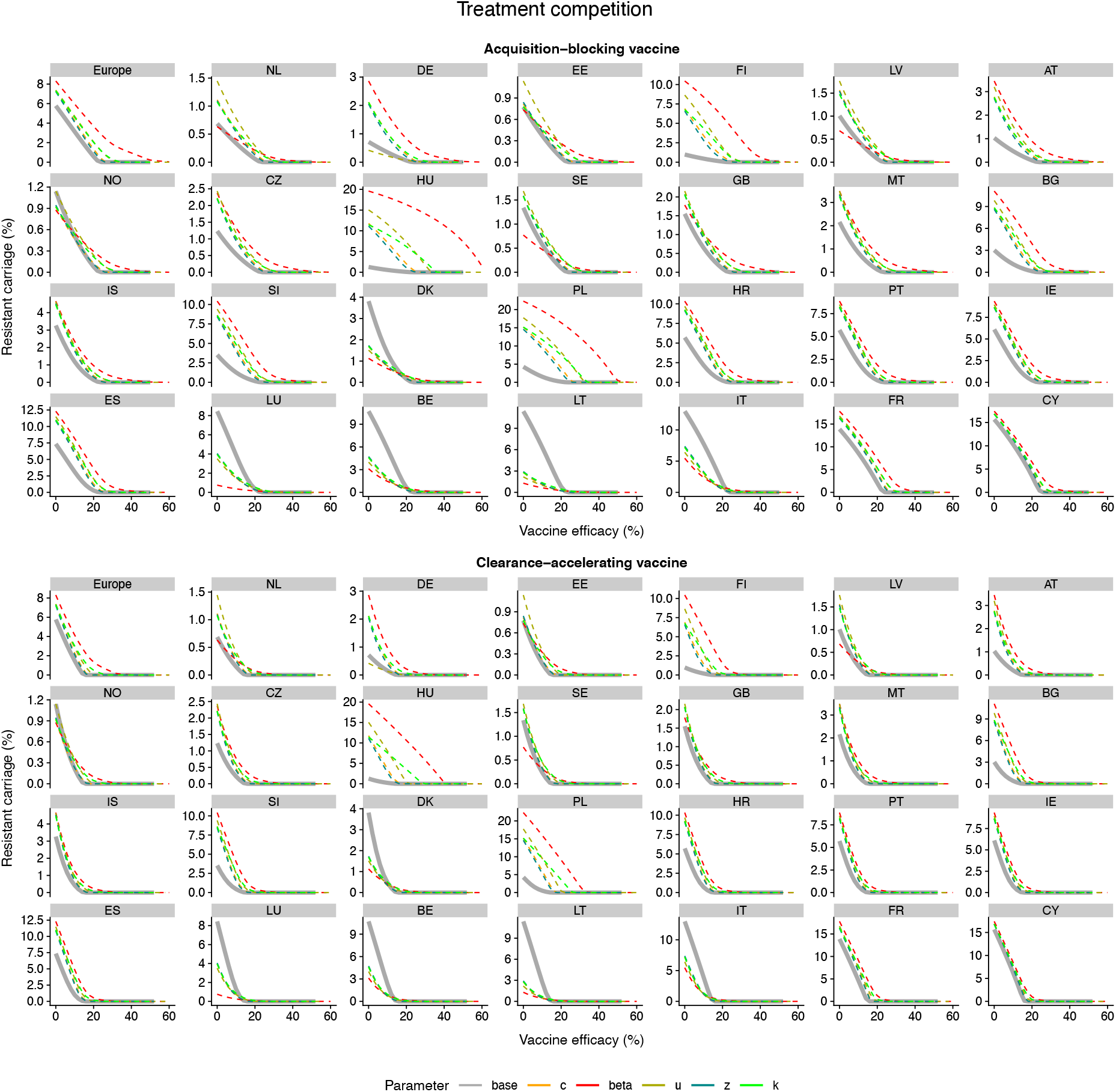
Vaccine impact for the “Treatment competition” model, varying parameters *β, c, u, κ*, and *z*. Impact of vaccination under the “Treatment competition” model, for those parameters able to capture the between-country variation in resistance frequency. The base model fit (thick grey solid line) is compared with the model fits in which parameters vary between countries (thin dashed lines).

**Fig. S12.**
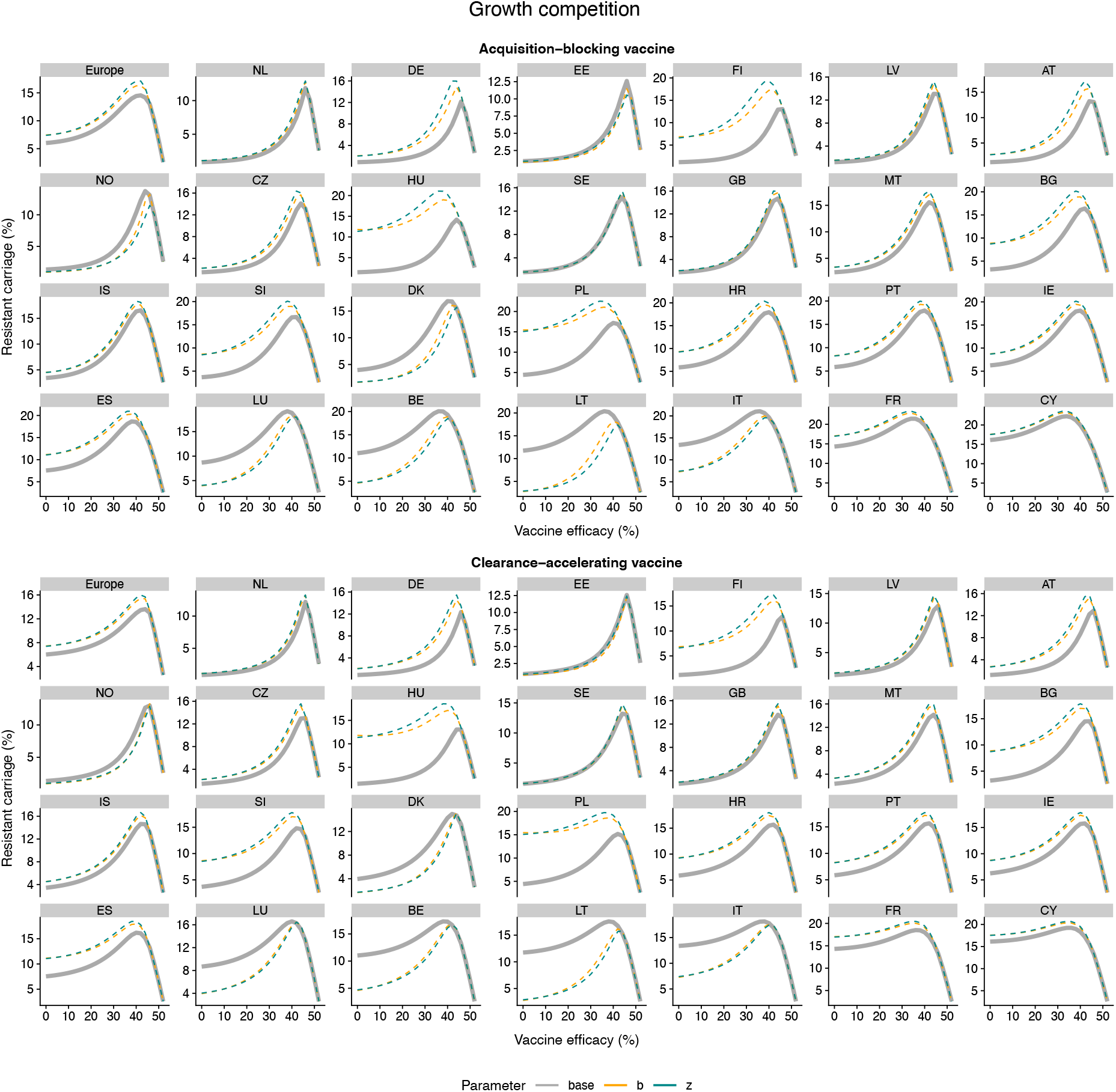
Vaccine impact for the “Growth competition” model, varying parameters *b* and *z*. Impact of vaccination under the “Growth competition” model, for those parameters able to capture the between-country variation in resistance frequency. The base model fit (thick grey solid line) is compared with the model fits in which parameters vary between countries (thin dashed lines).

**Fig. S13.**
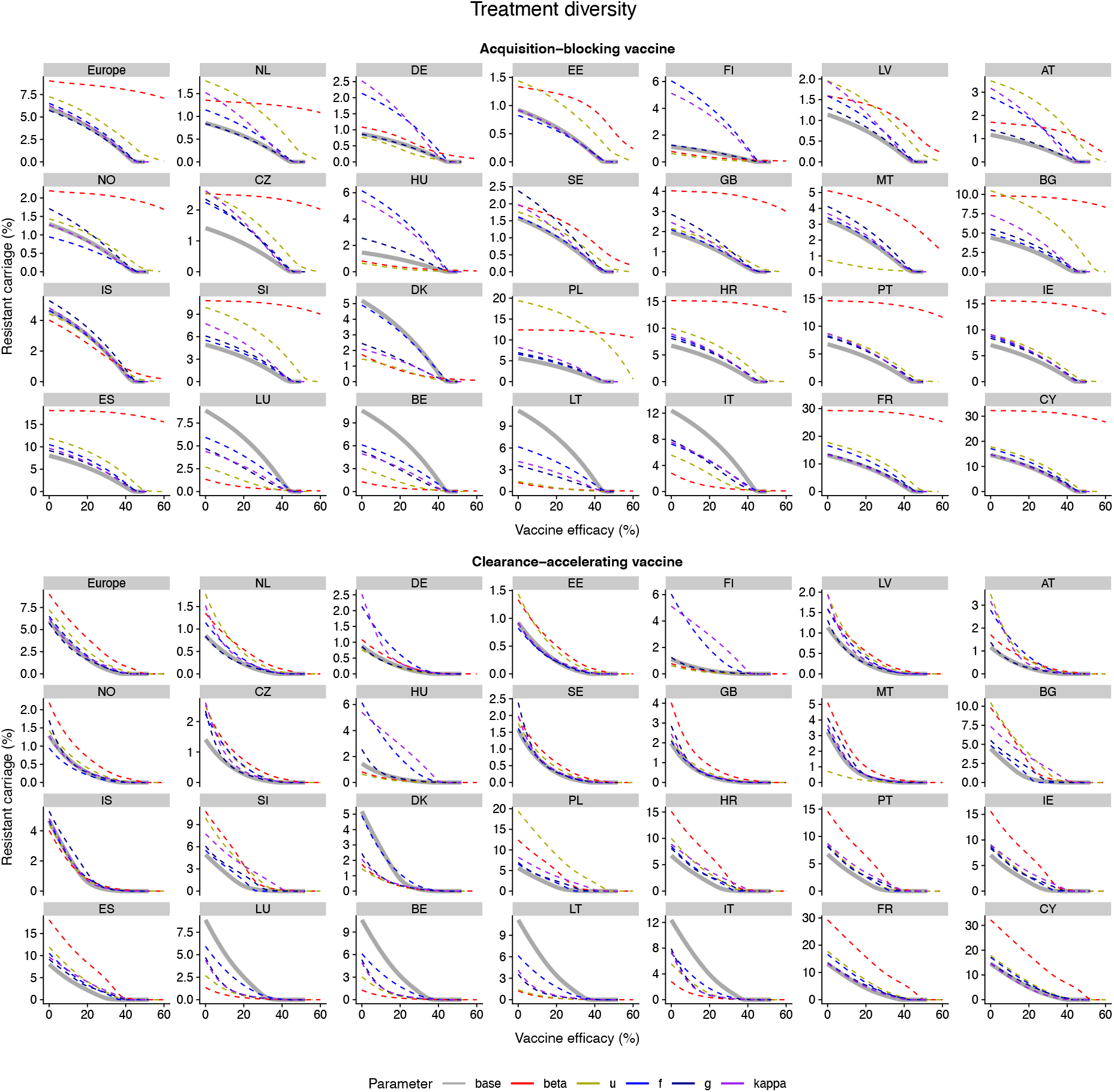
Vaccine impact for the “Treatment diversity” model, varying other parameters. Impact of vaccination under the “Treatment diversity” model, for those parameters *not* able to capture the between-country variation in resistance frequency. The base model fit (thick grey solid line) is compared with the model fits in which parameters vary between countries (thin dashed lines).

**Fig. S14.**
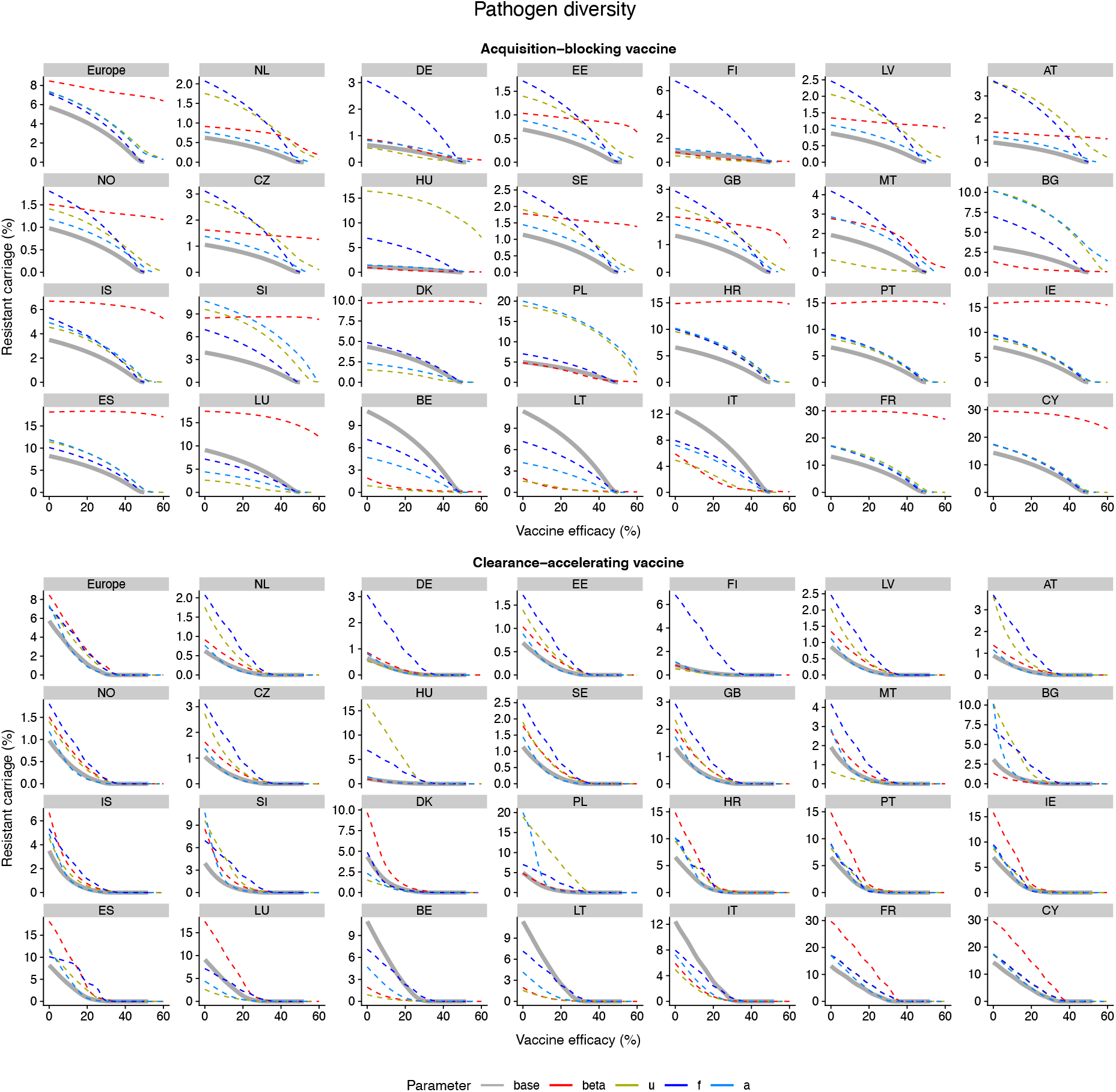
Vaccine impact for the “Pathogen diversity” model, varying other parameters. Impact of vaccination under the “Pathogen diversity” model, for those parameters *not* able to capture the between-country variation in resistance frequency. The base model fit (thick grey solid line) is compared with the model fits in which parameters vary between countries (thin dashed lines).

**Fig. S15.**
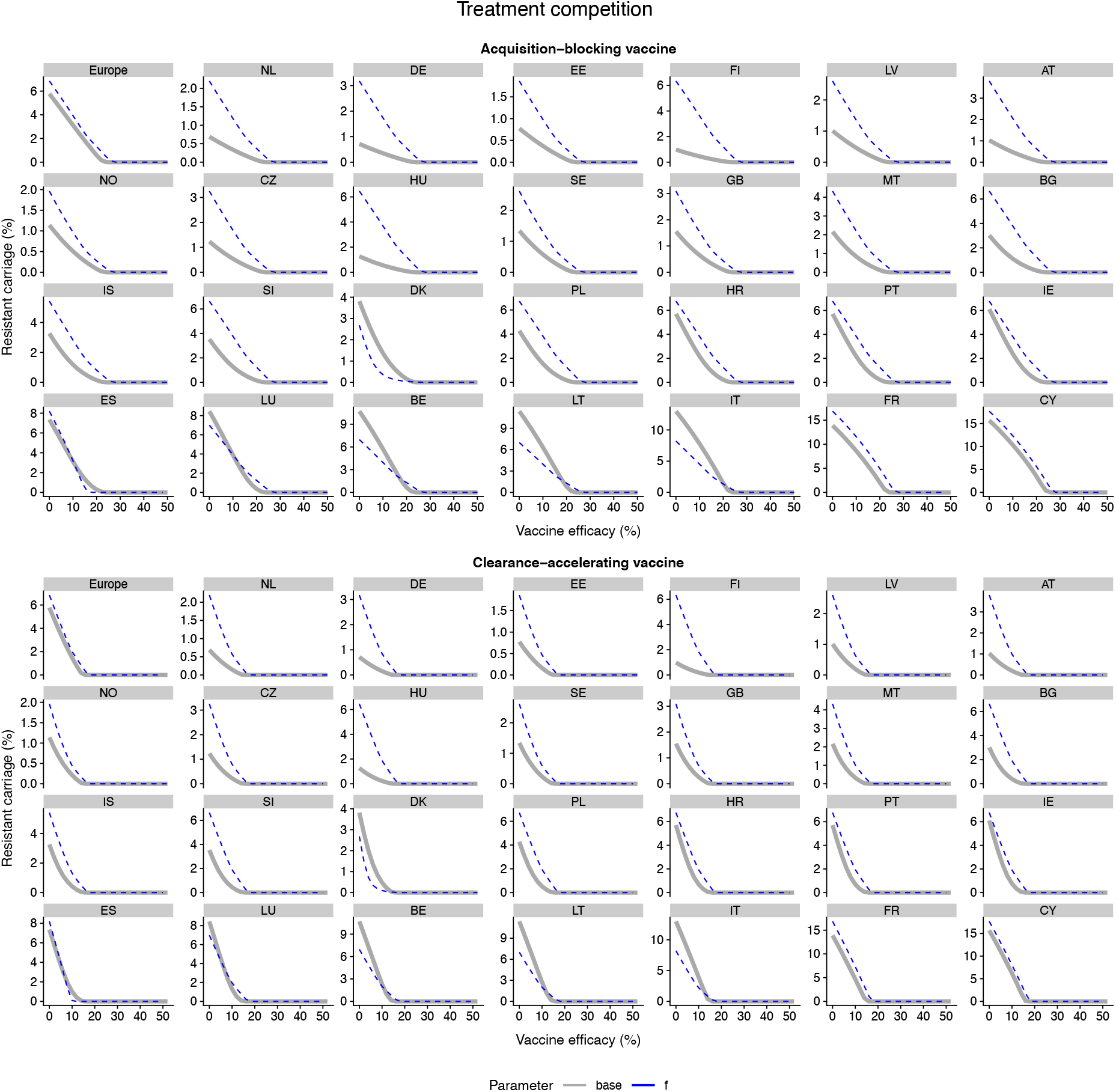
Vaccine impact for the “Treatment competition” model, varying other parameters. Impact of vaccination under the “Treatment competition” model, for those parameters *not* able to capture the between-country variation in resistance frequency. The base model fit (thick grey solid line) is compared with the model fits in which parameters vary between countries (thin dashed lines).

**Fig. S16.**
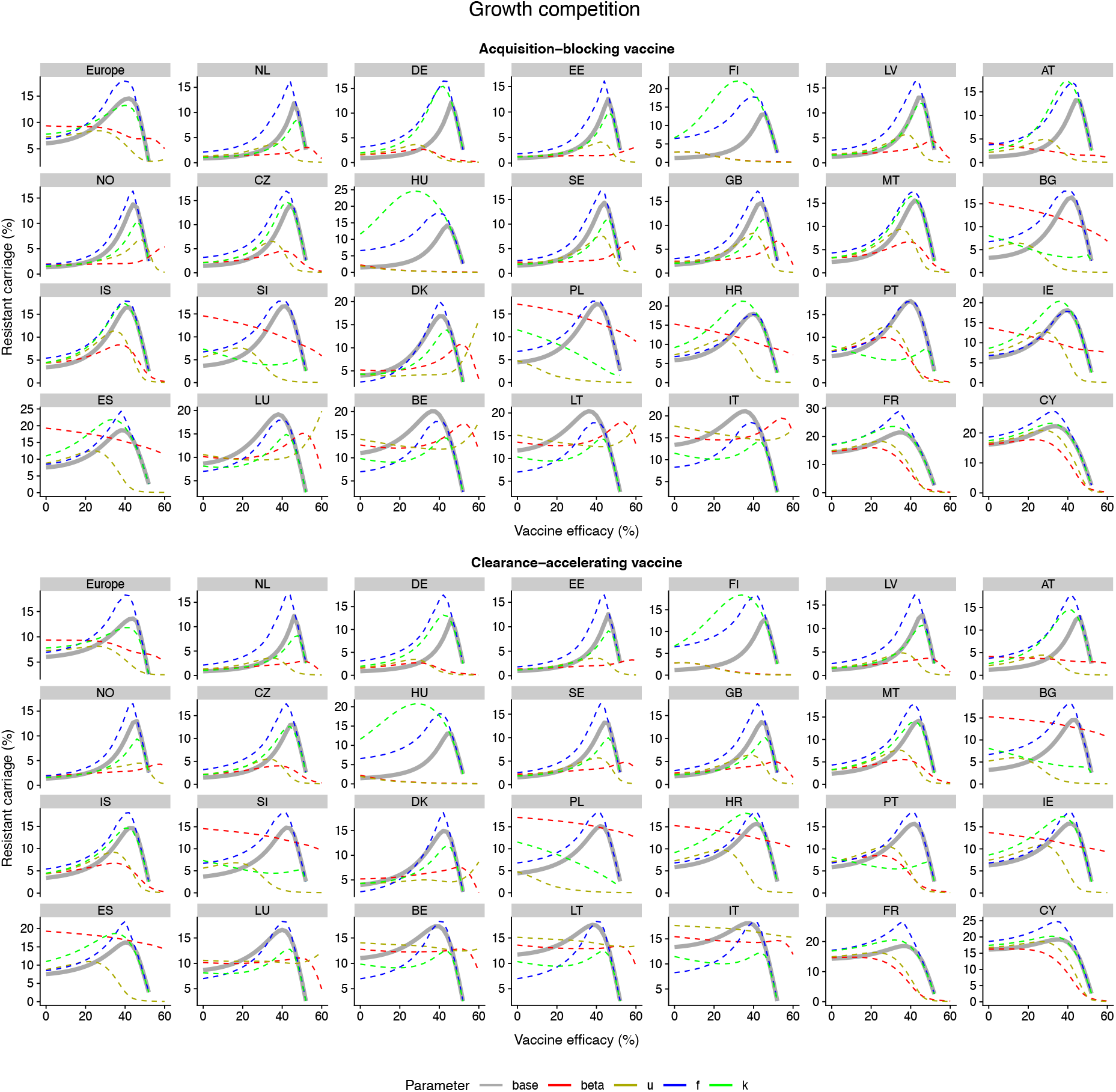
Vaccine impact for the “Growth competition” model, varying other parameters. Impact of vaccination under the “Growth competition” model, for those parameters *not* able to capture the between-country variation in resistance frequency. The base model fit (thick grey solid line) is compared with the model fits in which parameters vary between countries (thin dashed lines).

## Tables S1-S7

These tables can be found in an Excel spreadsheet accompanying the article.

*Table S1. Literature review —* Details of the literature review used to identify mechanisms for maintaining coexistence between sensitive and resistant bacterial strains.

*Table S2. Summary of model parameters —* Table describing model parameters and assumed values or prior distributions for model fitting.

*Table S3. Carriage duration —* Calculation of mean pneumococcal carriage duration for children under 5 years old in European settings.

*Table S4. Penicillin consumption —* Calculation of the mean number of defined daily doses of penicillin corresponding to a single treatment course for children under 5 years old in European countries.

*Table S5. Pneumococcal morbidity —* Calculation of the annual number of pneumococcal pneumonia cases in children under 5 in Europe and Kenya.

*Table S6. Carriage duration (Kilifi) —* Calculation of mean pneumococcal carriage duration for children under 5 years old in Kilifi, Kenya.

*Table S7. MCMC diagnostics —* Widely Applicable Information Criteria (WAIC), Leave-One-Out Information Criteria (LOOIC), effective posterior sample size and Gelman-Rubin diagnostics for Bayesian inference model fitting using Markov chain Monte Carlo.

